# Gut microbes contribute to variation in foraging intensity in the honey bee, *Apis mellifera*

**DOI:** 10.1101/2023.08.31.555606

**Authors:** Cassondra L. Vernier, Thi Lan Anh Nguyen, Tim Gernat, Amy Cash Ahmed, Zhenqing Chen, Gene E. Robinson

**Affiliations:** Carl R. Woese Institute for Genomic Biology, University of Illinois at Urbana-Champaign, Urbana, Illinois, USA; Department of Entomology, University of Illinois at Urbana-Champaign, Urbana, Illinois, USA; Neuroscience Program, University of Illinois at Urbana-Champaign, Urbana, Illinois, USA

**Author notes:** Corresponding Authors (CLV), (GER). Biology Department, Franciscan Missionaries of Our Lady University, Louisiana, USA.

## Abstract

Gut microbiomes are increasingly recognized for mediating diverse biological aspects of their hosts, including complex behavioral phenotypes. While many studies have reported that experimental disruptions to the gut microbiome result in atypical host behavior, studies that address how gut microbes contribute to adaptive behavioral trait variation are rare. Eusocial insects represent a powerful model to test this, due to their simple microbiomes and complex division of labor characterized by colony-level variation in behavioral phenotypes. While previous studies report correlational differences in gut microbiome associated with division of labor, here, we provide evidence that gut microbes play a causal role in defining differences in foraging behavior between honey bees. Gut microbial community structure consistently differed between hive-based nurse bees and bees that leave the hive to forage for floral resources. These differences were associated with variation in the abundance of individual microbes, including *Bifidobacterium asteroides, Bombilactobacillus mellis,* and *Lactobacillus melliventris*. Manipulations of colony demography and individual foraging experience suggested that differences in microbiome composition were associated with task experience. Moreover, single microbe inoculations with *B. asteroides*, *B. mellis,* and *L. melliventris* caused changes in foraging intensity. These results demonstrate that gut microbes contribute to division of labor in a social insect, and support a role of gut microbes in modulating host behavioral phenotypic variation.

## Introduction

Gut microbiomes are emerging as important drivers and modulators of host phenotype [1], with evidence supporting the role of microbiomes in host digestion and nutrition, immune health, development, and, more recently, behavior [2–7]. Over the past decade, the “microbiota-gut-brain axis” [7], which describes bidirectional interactions between the gut microbiome and host brain and behavior, has developed as an increasingly important field of study with implications for understanding the ecology and evolution of host-microbe interactions and animal behaviors. However, current studies of the microbiota-gut-brain axis across animal taxa are largely correlational, and causal studies exploring the relationship between gut microbes and host behavior are rare [7].

The functional relationship between hosts and microbiomes varies across host traits. In many cases, this relationship is obligatory, where the microbiome is necessary for normal functioning of the host [7–10]. Recently, it has been proposed that gut microbiomes play a facultative role in host phenotype, such that gut microbiomes contribute to phenotypic variation between individuals [11, 12]. Results in support of a facultative relationship come from studies across taxonomic groups. These include variations in non-behavioral and behavioral phenotypes [11, 12], such as behaviors associated with aging and senescence [13–18] and severity of neurodevelopmental disorders [19–21], as well as adaptive variations in behavioral traits both within [22, 23] and between species [24]. Eusocial insects represent a powerful model for understanding how gut microbes contribute to adaptive behavioral trait variation due to their relatively simple and stable gut microbiomes and their complex division of labor [25–27]. This division of labor is characterized by polyphenism between reproductive and non-reproductive individuals, as well as colony-level behavioral trait variation between non-reproductive individuals performing different tasks [25, 28]. In eusocial insects, gut microbes have been shown to contribute to natural variation in memory [22] and social interactions [29]. Likewise, studies indicate an association between gut microbiome and division of labor in eusocial insects [25, 30– 33]. However, whether gut microbes play a causal role in any aspect of eusocial insect division of labor remains unknown.

We use the European honey bee, *Apis mellifera,* to investigate the causal relationship between the gut microbiome and division of labor. Honey bees are a highly eusocial insect with well-characterized and tractable social behaviors and gut microbiomes. In typical honey bee colonies, sterile worker bees exhibit age-related division of labor, which is based on a pattern of individual behavioral maturation. Typically, adult worker bees perform brood care (“nursing”) and other activities in the hive during the first 1-3 weeks of adult life and then transition to foraging behaviors outside of the hive for the final 1-2 weeks of their life [28, 34]. However, honey bee division of labor is flexible and responsive to changing colony needs, and thus does not exclusively rely on the typical age patterns of behavioral maturation [35–39]. In addition, honey bee foragers exhibit considerable individual variation in preference for nectar or pollen [40, 41], exploratory behavior [42], and general foraging intensity [43]. Therefore, honey bee division of labor involves individual variation in both behavioral maturation and task performance [44].

The honey bee gut microbial community, most of which resides in the hindgut, is well-characterized and relatively simple, composed of ∼10-20 different species of facultatively anaerobic and microaerophilic host-adapted bacteria within the taxonomic groups of Actinomycetes (*Bifidobacterium*), Lactobacillaceae (*Bombilactobacillus* Firm-4 and *Lactobacillus* Firm-5), Gammaproteobacteria (*Gilliamella* and *Friscella*), Alphaproteobacteria (*Bartonella* and *Bombella*), and Betaproteobacteria (*Snodgrassella*) [27]. Furthermore, the individual members of the honey bee gut microbiome are consistently present, but differ in abundance across different individuals and populations of honey bees [27]. Of particular interest, the composition of honey bee gut microbial communities can be experimentally manipulated. All species can be cultured [45] and used to inoculate young bees [29, 46, 47], who must acquire their microbiome from older bees or hive materials [48]. Furthermore, previous studies indicate an association between the gut microbiome and various aspects of division of labor in the honey bee [30–32, 49, 50]. Due to these attributes of the honey bee gut microbiome, it is a great model for understanding the causal effects of gut microbes on complex host behaviors. We took advantage of these behavioral and microbiome features to determine whether gut microbes play a causal role in honey bee division of labor, with a specific focus on behavioral maturation and variation in foraging intensity.

## Materials and Methods

### Animal Husbandry, treatments, and collections

Honey bee colonies were managed using standard beekeeping techniques at the University of Illinois Bee Research Facility in Urbana, IL. Honey bees in this area are a genetic mixture of subspecies, primarily *Apis mellifera ligustica* and *carnica* subspecies. For collection of nurses and foragers, nurses were identified as those actively feeding brood on a brood frame, and foragers were identified as those returning to the hive with pollen loads on their hind legs or having a distended abdomen due to nectar loads [37, 51]. To reduce genetic variation between workers in the colonies used for the single-cohort colony, big-back colony and single microbe inoculation studies (described below), we used bees derived from queens that were each instrumentally inseminated with sperm from a different single drone (SDI) (queen rearing and inseminations were performed by Sue Cobey, Honey Bee Insemination Service, Washington State University and Dr. Osman Kaftanoglu, Apimaye USA). Sample size for 16S rRNA sequencing studies was 10 bees per group as in [52]. All bees used in 16S rRNA sequencing analyses were washed once with 12.5% bleach in water and twice with double deionized water and flash frozen. All samples were stored at -80°C until further analysis.

### Single- and double-cohort colonies

Single-cohort colonies (SCC) were established in the summer of 2020 as previously described [36, 37, 51]. Honeycomb frames with pupal brood were taken from one or more source colonies derived from the above-mentioned queens and placed in a 34°C incubator at 60-65% relative humidity. About 1000 newly eclosed bees (<24 hours old) were gently brushed from these frames <24 hours later and placed in a small five-frame hive box with a new, unrelated mated queen, one honey frame, an empty honeycomb frame, and three new frames with wax-covered plastic foundation upon which bees can build new wax honeycomb. SCCs were kept outdoors, and bees were collected as typical-age nurses and precocious foragers at approximately one week of age (7-9 days of age, variation due to rainy weather), and as over-age nurses and typical-age foragers at three weeks of age. To obtain over-age nurses, frames containing brood were removed and replaced with empty comb two weeks after queen introduction to ensure that a new cohort of bees did not eclose before collections. For the SCC depicted in Fig. 2 and Supplementary Fig. 1, bees from four distinct SDI source colonies (unrelated queens and drones) were used to construct a single SCC. Four SDI source colonies were used to obtain the necessary 1000 newly eclosed bees, and allowed us to test for the role of source colony in gut microbiome development (Supplementary Fig. 1). For each collection (Day 9 and Day 21), 10 bees of each source colony were collected for each behavioral task group. On Day 9, nurses were only able to be collected for two source colonies. For the SCC depicted in Supplementary Fig. 4, bees from a single SDI source colony were used, and groups of 10 bees were collected at each collection (Day 7 and Day 21) for each behavioral task group.

Double-cohort colonies (DCC) were used to assess the effects of different inoculation treatments (see below) on foraging behaviors in a common colony environment. DCCs were established similarly to SCCs with a new, mated queen, 1 plastic honeycomb frame provisioned with honey and pollen paste, 1 empty plastic honeycomb frame, ∼200 7-day old barcoded treatment bees (100 of bees from a single microbe inoculated treatment group and 100 from a corresponding microbiota-depleted group) and ∼800 newly eclosed unbarcoded “background” bees (mixed colony backgrounds), whose addition contributed to accelerated behavioral maturation of the older cohort [51, 53]. In the presence of the older cohort, younger “background” bees are not expected to contribute to the foraging force [54] and did not throughout the course of the experiment.

### Big-back colonies

Big-back colonies were established as previously described [36, 51, 53] and kept outdoors in the summer of 2021. For each colony, ∼800-900 newly eclosed bees from a single SDI source colony were used; half with a paint mark on the thorax, and half with a plastic tag attached to the thorax (∼3 mm diameter, ∼1 mm thick; “big-back” bees). Four days later, ∼400 newly eclosed bees from ∼10 mixed source colonies (headed by naturally mated queens) were introduced to increase the proportion of precocious foragers in the first cohort [36, 51, 53]. The entrance to the hive was blocked by a piece of Plexiglas with holes in it to prevent the big-back bees from leaving the hive, but to allow paint-marked bees to come and go freely. We observed that guard bees patrolled the hive entrance on the inside of the Plexiglas piece.

Bees were collected at 10 days of age: returning active foragers and nurses collected as described above, and big-back/inactive forager bees collected as they were attempting to leave the hive via the holes in the plastic, indicating they too were in the behavioral state of foraging but had never left the hive. 10 bees per group were collected from all three colonies, with the exception that only seven nurses from the focal group in colony 2 were found and collected.

### Gut microbiome DNA extraction, 16S rRNA sequencing and analysis

Frozen honey bee guts were dissected under sterile conditions on dry ice. Combined mid- and hind-guts of individual bees were homogenized by maceration with a disposable sterile pestle (VWR, Radnor, PA, USA). The homogenate was added to a PowerSoil Pro Bead Solution tube (Qiagen, Germantown, MD, USA), and DNA was extracted using a DNeasy PowerSoil Pro DNA isolation kit (Qiagen), following manufacturer’s instructions. The hypervariable V4 region of the 16S rRNA gene was amplified by PCR in triplicates, with a negative control (no DNA), using Platinum Hot Start PCR Master Mix (Invitrogen, Waltham, MA, USA), primers and barcodes designed in [55] with a final concentration 0.25 μM, and the following cycling conditions: 94° 3 min, 35x[94° 45s/50° 60s/72° 90s], 72° 10 min. Before sequencing, PCR products were visualized on agarose gels to confirm negative controls did not have amplification, while all samples had expected amplification. All samples met these criteria, indicating no contamination. Samples were pooled based upon concentrations and sequenced on an Illumina MiSeq with 2×250bp paired-end reads. The samples were split among three sequencing runs: #1: SCCs (Dec 2020), #2: Big-back colonies (July 2021), #3: Typical colony nurses and foragers, single microbe inoculated bees (Dec 2021). Reads from these sequencing runs can be found in NCBI’s Sequence Read Archive PRJNA958253. Sequences for typical colonies from naturally mated queens were retained from [49].

Samples were demultiplexed using QIIME2, and paired end reads were truncated at the first base with a quality score of <Q3 using DADA2 [56]. Paired-end reads were merged and amplicon sequence variants (ASV) were identified using DADA2 [57]. Chimeric ASVs were removed and the remaining ASVs were taxonomically classified using the BEExact database [58]. ASVs that were taxonomically identified as a bee specific genus by the BEExact database, but were unclassified at the species level, were subsequently classified to species level if possible, using NCBI megaBLAST. To do this, representative 16S rRNA gene V4 sequences for each ASV obtained from the QIIME2 rep-seqs.qza object were BLASTed against the entire NCBI nucleotide database, and unclassified ASVs were called as the first identified full species with the lowest E-score, if query cover > 80% and percent identity > 92%. Prior to statistical analyses, ASVs that were identified as mitochondrial or chloroplast were removed from the data. Mitochondrial and chloroplast reads were retained for visualization of single microbe inoculated and microbiota-depleted bee gut microbiomes.

For sequencing run #1, 3,220,034 (1,610,017 pairs) total sequence reads were obtained, 1,431,503 pairs were filtered, merged and identified as non-chimeric (90%), and 627 ASVs were identified. For sequencing run #2, 945,310 (472,655 pairs) total sequence reads were obtained, 424,785 pairs were filtered, denoised, merged and identified as non-chimeric (90%), and 110 ASVs were identified. For sequencing run #3, 2,538,722 (1,269,361 pairs) total sequence reads were obtained, 1,141,046 pairs were filtered, denoised, merged and identified as non-chimeric (90%), and 832 ASVs were identified.

To estimate the abundance of individual honey bee-associated microbial species in each sample, the read counts for all ASVs that matched the same species were combined. We retained the following honey bee-associated species: Acetobacteraceae related to *Commensalibacter* [possibly Alpha-2.1 and Alpha-2.2], *Bartonella spp.*, *Bifidobacterium asteroides*, *Bifidobacterium coryneforme* [syn. *Bifidobacterium indicum*], *Bombella apis* [previously *Parasaccharibacter apium*], *Frischella perrara*, *Gilliamella apicola*, *Lactobacillus apis*, *Lactobacillus helsingborgensis*, *Lactobacillus kullabergensis*, *Bombilactobacillus mellifer* [previously *Lactobacillus mellifer*], *Bombilactobacillus mellis* [previously *Lactobacillus mellis*], *Lactobacillus melliventris*, *Apilactobacillus kunkeei*, and *Snodgrassella alvi* [27, 59, 60]. We also retained other unclassified *Lactobacillus* species, *Limosilactobacillus spp.*, *Arsenophonus spp.*, *Klebsiella spp.*, and *Mixta spp.* because they made up a significant portion of reads (>500 reads in at least 1 sample). In the case of single-microbe inoculated and microbiota-depleted bees (see below), *Enterococcus spp.*, *Staphylococcus spp.,* and Unassigned (not able to be classified below Kingdom of Bacteria) were additionally retained for visualization. All other ASVs, each comprising 0-1% of total reads per sample, and representing microbes that were not classified in the aforementioned groups and/or are not typically associated with honey bees, were combined and labeled as “other.” The variation between samples in the abundance of “other” microbes is similar to previously published datasets [29, 31]. To calculate the proportion of each species in each sample, that species’ read count was divided by the total read count for each sample. Raw read counts and/or proportion data were used in analyses as described below. For the depiction of single microbe inoculated bee gut microbiomes, “other” ASVs were combined based on genus (or the lowest taxonomic level identified), and those representing >500 reads in at least 1 sample are depicted in a stacked barplot, while all genera are included in supplementary data on Mendeley Data (DOI: 10.17632/f2s47y3nhn).

Due to the compositional nature of 16S rRNA sequencing data, typical measures of beta diversity and differential abundance analyses have limitations [61–63]. Therefore we followed analyses outlined in [61]. In short, for beta-diversity analyses, we used clr-transformations of raw read counts for each sample in statistical tests and Aitchison distance for visualization (see “Statistical analysis” section below). To estimate the differential relative abundance of individual microbial species between samples through tools that have a compositional foundation (referred to as “relative abundance” throughout the text), we used Analysis of Composition of Microbiomes with Bias Correction (ANCOM-BC) [62, 64, 65] on raw read counts. This test relies on the total sample read count and relative abundance of individual microbes in each sample to estimate the true abundance of individual microbes in each gut ecosystem while accounting for the compositional nature of sequencing data [64].

To estimate absolute bacterial species abundances in individual samples (“absolute abundance”), we quantified the bacterial load in each sample using quantitative PCR (qPCR), as in [29, 30]. We performed standard curves using serial dilutions of plasmids (TOPO pCR2.1 (Invitrogen)) containing the target sequence (10^8^-10^3^ copies per 3 μl), which were calculated from the molecular weight of the plasmid and the DNA concentration of the plasmid. Primer efficiencies were measured using: E = 10^(-1/slope)^ [66], and the copy number of each target in 3 μl of DNA sample was calculated from the sample’s Ct score, primer efficiency, and standard curve using: n = E^(intercept-Ct)^x(DNA extraction elution volume/3) [29, 30]. These values were calculated for both the 16S rRNA gene and the *Actin* gene. To account for differences in DNA extraction efficiency, we calculated a normalized number of 16S rRNA gene copies by dividing the number of 16S rRNA gene copies by that sample’s number of *Actin* gene copies and multiplying this by the median number of *Actin* copies for that experiment [29, 30]. To calculate the absolute abundance of each microbial species in each sample, the relative abundance (proportion) of each species in each sample was multiplied by the normalized number of 16S rRNA gene copies in that sample [29, 30]. We then took the log10 value of each of these numbers and used these in further analyses, replacing 0s with 1s before we did so.

PCRs were performed in triplicate using 10 μl reactions of 0.5 μM primers targeting the 16S rRNA gene (F: AGGATTAGATACCCTGGTAGTCC, R: YCGTACTCCCCAGGCGG) or the *A. mellifera Actin* gene (F: TGCCAACACTGTCCTTTCTG, R: AGAATTGACCCACCAATCCA) [46] in 1X *Power*SYBR Green PCR Master Mix (Applied Biosystems, Waltham, MA, USA) with 3 μl of DNA. Each primer set included a no template control. Reactions were performed in 384 well plates using a QuantStudio 6 Flex (Applied Biosystems) with the following cycling conditions: 50° 2 min, 95° 10 min, 40x[95° 15 sec/60° 1 min], melt curve: 95° 15 sec, 60° 1 min, 95° 15 sec. Standard curves using serial dilutions of plasmids (TOPO pCR2.1 (Invitrogen)) containing the target sequence (10^8^-10^3^ copies per 3 μl) were calculated from the molecular weight of the plasmid and the DNA concentration of the plasmid.

We found an effect of colony replicate on gut microbial community structure across all experiments, indicating that bees from different colonies have different gut microbial communities and that early environment and/or genetics influences the composition of the gut microbiome. This was the case when comparing bees from different typical colonies (Supplementary Fig. 1A- B), among SCC bees originating from different colony sources (Supplementary Fig. 1C-D), and across bees from different big-back colonies (Supplementary Fig. 1E). Colony differences in honey bee microbiome composition have been previously reported [52].

### Single microbe inoculations

To identify effects of individual microbes on behavior we treated groups of newly eclosed bees, who emerged under sterile lab conditions, with either an inoculum of a microbe of interest or sterile food in order to produce bees whose microbiomes were composed of a single honey bee-associated microbe (single microbe-inoculated) or no honey bee-associated microbes (“microbiota-depleted,” historically referred to in this way since bees lacking a typical honey bee microbiome are not completely microbe-free [46, 67]), respectively. Although single microbe inoculations do not represent a natural microbiome, we chose this approach because it would likely allow for the most control over microbiome composition across replicates. This is because inoculations with multiple microbes may result in variation in microbiome structure between individuals, groups, and/or replicates since honey bee microbiome structure is shaped by both environmental and genetic factors [27, 48, 68]. These experiments occurred in the summer of 2021.

To obtain bees that lack the dominant honey bee bacterial species in the gut, modified methods from [46, 67] were used. Specifically, tan-colored pupae with dark eyes were gently removed from brood frames from three or four source colonies per trial using ethanol sterilized forceps, placed dorsal side down in sterilized 3D-printed dental grade resin modular pupation plates (Supplementary Fig. 2, design file on Mendeley Data), which were fitted into sterilized Plexiglas cages, and kept in a 34°C incubator with 60-65% relative humidity for 2 days [69]. Eclosed bees were moved in groups of 30-40 to sterilized Plexiglas boxes (10 x 10 x 7 cm) [70] in a 32°C incubator at 60-65% relative humidity (5 boxes per treatment for each replicate). Each box was provisioned with a ∼1 g sterilized pollen patty and 1.7 mL autoclave (15 minutes) sterilized 25% sucrose with or without inoculum. An equal number of bees from each source colony were added to each box, such that each treatment group (microbiota-depleted or single microbe inoculated) had an equal number of bees of each colony background.

For single microbe inoculations, microbes were obtained from the DSMZ Leibniz-Institut in Germany (*B. asteroides* DSM 20089, *B. mellis* DSM 26255, *L. melliventris* DSM 26256) and were streaked on solid media and then cultured under anaerobic conditions at 35°C for 3 nights in de Mann, Rogosa, Sharpe (MRS) (*B. asteroides*) or MRS + 20g/L fructose (*B. mellis and L. melliventris*) broth, and glycerol stocks were made and kept at -80°C until future use. Ten days prior to the onset of inoculation experiments, glycerol stocks were streaked on solid MRS (*B.asteroides*) or MRS + 20g/L fructose (*B. mellis* and *L. melliventris*) media plates and were grown under anaerobic conditions at 35°C for 3 nights (Oxoid Anaerojar with Anaerogen bags, Thermo Scientific, Waltham, MA, USA). A single colony of bacteria from each plate was cultured under anaerobic conditions (Anaerobic Hungate Culture Tubes with air displaced by CO2, VWR, Radnor, PA, USA), shaking at 35°C for 3 nights and was then retained at 4°C. Every day, starting 2 days prior to the onset of inoculations, new cultures from this same original stock were prepared by culturing 50 μl of the stock in 5 mL of fresh MRS/MRS + 20g/L fructose broth under anaerobic conditions, shaking at 35°C for 2 nights. These cultures were spun down (3000 rpm for 5 min) and resuspended in 1x PBS to an OD of 1 and 50 μl of a new, fresh culture solution was added to 1.7 mL sterile 25% sucrose water in a new inverted microtube for each treatment box every morning over the course of a 5-day inoculation period. For microbiota-depleted bees, 50 μl of sterile 1x PBS was added to 1.7 mL 25% sucrose water in a new inverted microtube for each treatment box every morning over the course of a 5-day inoculation period. Bees were then fed sterile 50% sucrose water for 2 days following the 5-day inoculation period. Very few bees (1-2 per box) died during this inoculation period. Bees were kept in treatment boxes until 7 days old. Unfortunately, we did not keep track of the treatment box each bee came from, rather bees for each treatment were pooled and randomly chosen for 16S rRNA analysis or behavioral assays. Microbe survival in 25% sucrose water was ensured by placing 50 μl of each microbial culture (OD ∼1) in 1.5 mL 25% sucrose water overnight and then plating this solution using species specific culturing conditions.

### Foraging assays and analysis using barcoded bees

To obtain comprehensive records of foraging activity, we used an automated behavioral tracking system (“bCode”) that uses a custom matrix barcode, enabling the unique identification of individual bees [71]. After single microbe inoculations, 7-day old inoculated and microbiota-depleted bees were cold-anaesthetized, a unique barcode was affixed to their thorax using super glue, and a spot of colored paint associated with their treatment group was placed on their abdomen. An equal number (∼100) of bees from a single microbe inoculated group and a corresponding microbiota-depleted group (same mix of colony backgrounds, same age) were given barcodes from different sets, and were placed together in a DCC. Four colony replicates per microbe treatment were performed. Two experimental days, corresponding with *B. mellis* colony replicate 3 days 1-2 and *L. melliventris* colony replicate 2 days 1-2, experienced bad weather, and thus no foraging occurred on those days.

Colonies were kept in a dark, temperature (32°C) and humidity (∼50%) controlled building, with access to the outside environment through a plastic tube. To monitor flight activity, an entrance monitor with a video camera (Supplementary Methods) was attached to the hive entrance, on the outside-facing end of this tube (Supplementary Fig. 3). The camera recorded videos of bees entering and leaving the hive from 05:00 to 21:00 daily for a total of six days. Barcodes in these videos were detected as in [71], and detections were filtered to remove tracking errors and misidentifications. A mean of 97.82%, with a range between 96.24%-99.02%, of detections were retained after these filtering steps. We then applied a flight activity detector [72, 73] (Supplementary Methods) to the remaining barcode detections to identify passes through the entrance monitor. Computer code for this detector can be found at: https://github.com/gernat/btools.

Passes through the entrance monitor determined by the flight activity detector were filtered to remove unused barcodes, whose detections either occurred through detection error or due to bees accidentally entering the wrong experimental colony. A mean of 99.19%, with a range of 95.4%-100%, of events were retained after this filtering. Passes were then classified as “incoming,” “outgoing,” or “other.” Since incoming passes had a lower error rate than outgoing passes (Supplementary Table 8), incoming passes alone were used to denote a foraging trip. Individual foraging trips that occurred within 5 min of each other were condensed into a single event. A bee’s first day of foraging was defined as the first day on which it performed at least 4 foraging trips, with at least 50% of these trips occurring during peak foraging hours (11:00-15:00 CST), as per previous studies [43, 74]. These criteria yield similar automated thresholds corresponding to human observations of foraging behaviors [43]. All incoming passes from this first day of foraging were counted as foraging trips, as were all subsequent incoming passes [43]. Likewise, an individual bee was considered a “forager” from this day forward. Due to the large size of these data sets (owing to high dimensional automated behavioral monitoring) the number of foraging trips per bee and treatment group and the number of foragers per treatment group on each experimental day are available on Mendeley Data (DOI: 10.17632/f2s47y3nhn).

To quantify the degree of skew in foraging intensity among all workers, we calculated the Gini coefficient [43, 75] for each experimental colony across all days and each experimental colony on each day, since colony dynamics change with changes in weather and resource availability, and thus likely changed throughout our studies. The values we obtained (Table 2) are similar to those in previous studies [43, 75], indicating differences in foraging intensity between individuals in each colony. To determine which bees performed the majority of the foraging for each experimental colony on each day, we ranked individuals based on the proportion of total foraging trips they performed for the colony on each day. A group of “elite foragers” was defined as the subset of bees (variation, 5-40% of the foragers) performing ≥50% of the foraging trips for each colony on each day [43, 75]. Number of elite foragers per treatment group on each day are available on Mendeley Data (DOI: 10.17632/f2s47y3nhn).[74]

### Statistical Analysis

All statistical analyses were performed in R (v 4.2.0) [76]. For all analyses, assumptions (e.g., normality, homogeneity of variances) were checked before statistical analysis. The following functions were used in the base statistics package: “shapiro.test” (Shapiro-Wilk Normality Test), “fligner.test” (Fligner-Killeen Test of Homogeneity of Variances), “t.test” (Student’s t-test), “wilcox.test” (Mann-Whitney Rank Sum Test), “aov” (ANOVA), “TukeyHSD” (Tukey’s HSD post hoc test), “Kruskal.test” (Kruskal-Wallis test). Kruskal-Wallis post hoc was performed using “kwAllPairsDunnTest”, PMCMRplus package [77]. Microbiome beta diversity was analyzed using Permutation MANOVAs with 999 permutations (“adonis2”, vegan package [78]) on clr-transformed (“transform”, microbiome package [79]) raw read counts, followed by Pairwise Permutation MANOVAs with 999 permutations and FDR p-value adjustment (“pairwise.adonis”, pairwiseAdonis package [80]), and visualized using Principal Components Analysis (“ordinate”, phyloseq package [81]) with Aitchison distance (“distance”, phyloseq package [81]). The relative abundance of individual microbes was analyzed using raw read counts and ANCOM-BC with FDR adjustment (“ancombc2”, ANCOMBC package [62, 65]) with task as a fixed effect and colony as random effect. Relative abundances were visualized using stacked barplots (“ggplot”, ggplot2 package [82]). To compare the absolute abundance of individual microbes, Permutation ANOVAs were used with 999 permutations (“perm.anova,” RVAideMemoire package [83]), and p-values were adjusted for multiple comparisons (“p.adjust” with FDR adjustment). Absolute abundance data were visualized using dotplots (“ggplot,” ggplot2 package [82]). Age at onset of foraging data (using each individual bee’s first day of foraging) were analyzed using a Cox Proportional Hazards model (“coxph,” survival package [84]), stratified by replicate [35, 74], and depicted as survival plots using the cumulative proportion of bees from each treatment group that were identified as foragers on each day. Proportions of foraging trips per individual and last day of foraging data were checked for outliers (“identify_outliers,” rstatix package [85]) and were analyzed as generalized linear mixed-effects models (“glmer,” lme4 package [86]) with log-normal distributions and nAGQ=0, inoculation treatment and day as main factors, and replicate and individual as random factors, followed by “Anova” (car package [87]) to determine main factor significance, and “emmeans” (emmeans package [88]) with Tukey’s p-value adjustment for pairwise comparisons. Linear mixed-effects models (“lmer,” lme4 package [86]) with inoculation treatment and day as main factors, and replicate as random factors, followed by “Anova” (car package [87]) to determine main factor significance, and “emmeans” (emmeans package [88]) with Tukey’s p-value adjustment for pairwise comparisons were used in the remaining behavioral analyses. Gini coefficients were calculated using “Gini” (DescTools package [89]). All behavioral data, except age at onset of foraging, were visualized using boxplots (“ggplot,” ggplot2 package [82]).

## Results

### Nurses and foragers differ in gut microbial community composition

To begin to investigate whether gut microbes influence honey bee division of labor, we compared gut microbial communities in nurse and forager bees, which represent two of the canonical behavioral task groups in honey bee division of labor, as well as two discrete time points in behavioral maturation [28]. Nurses and foragers differ markedly in physiology [37, 90–95], neuroanatomy [96], neurochemistry [42, 97, 98], gene expression [38, 99] and gene regulation [74, 100–102]. Therefore, we reasoned that testing for consistent differences in gut microbial community between these two groups would be a powerful way to test for associations between gut microbiome variation and division of labor.

We performed 16S rRNA sequencing on gut samples from nurses and foragers from three unrelated honey bee colonies and reanalyzed, using updated methods [56, 57], a similar dataset from a previously published study that used a different 16S rRNA sequencing method [49]. We found that nurses and foragers differed significantly in gut microbial community structure in both the previously published dataset (Fig. 1A) and our new dataset (Fig. 1B), indicating robust differences associated with division of labor and independent of sequencing method.

**Figure 1.**
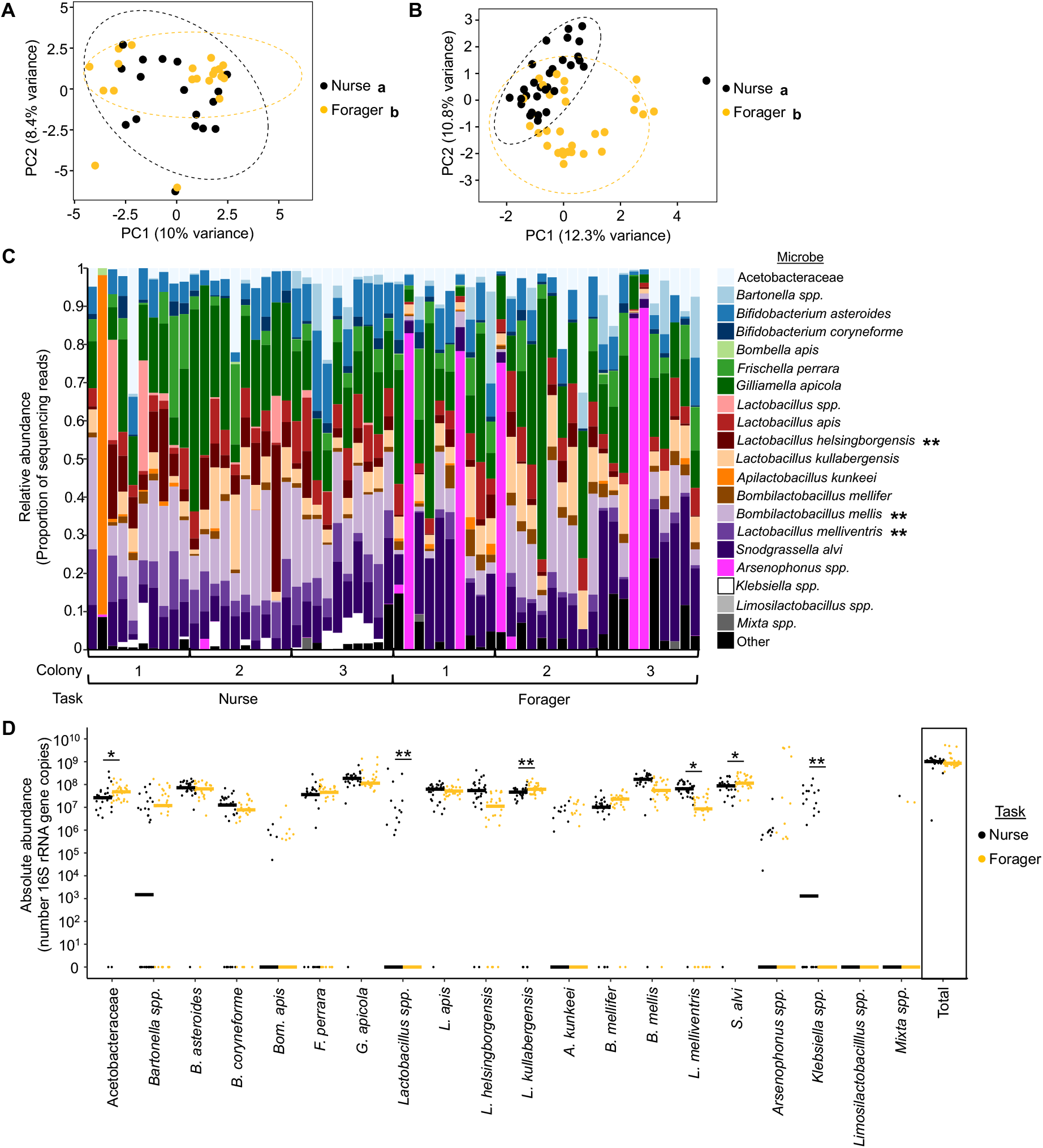
Nurses and foragers from typical honey bee colonies differ in gut microbial community. **(A-B)**Nurses and foragers differed in overall gut microbial community structure. (A) Reanalyzed data from Kapheim et al 2015. Two-way Permutation MANOVA using Aitchison Distance, Task: F1,38 = 1.51, R^2^ = 0.036, p = 0.016; Colony: F4,38 = 1.72, R^2^ = 0.165, p = 0.001, Task*Colony: F2,38 = 1.17, R^2^ = 0.056, p = 0.100. n = 1-12 bees/colony, 5 colonies. (B) New data. Two-way Permutation MANOVA using Aitchison Distance, Task: F1,59 = 4.28, R^2^ = 0.066, p = 0.001; Colony: F2,59 = 2.31, R^2^ = 0.071, p = 0.001, Task*Colony: F2,59 = 1.20, R^2^ = 0.037, p = 0.144. n = 10 bees/colony, 3 colonies. Depicted as Principal Components Analysis (PCA) plots. Lowercase letters in legends denote statistically significant groups. **(C)** Nurses and foragers differed in relative abundance of four individual microbial species (new data only). Depicted as stacked bar plots, with each bar representing a single bee’s microbiome. Asterisks in legend: *, pσ0.05, **, pσ0.01, ANCOM-BC between nurses and foragers. See Supplementary Table 1 for all p-values. **(D)** Nurses and foragers differed in absolute abundance of four individual microbial species but not in the total normalized number of 16S rRNA gene copies (new data only). 10^x^ number of 16S rRNA gene copies, calculated by multiplying the relative abundance each microbe in each sample (determined through 16S rRNA sequencing) by the normalized number of 16S rRNA gene copies in the sample (determined through qPCR). Depicted as dot plots with all data points plotted, line represents median, n = 10 bees/colony, 3 colonies. *, pσ0.05, **, pσ0.01, Permutation ANOVA Test between nurses and foragers. See Supplementary Table 1 for all p- values.

**Figure 2.**
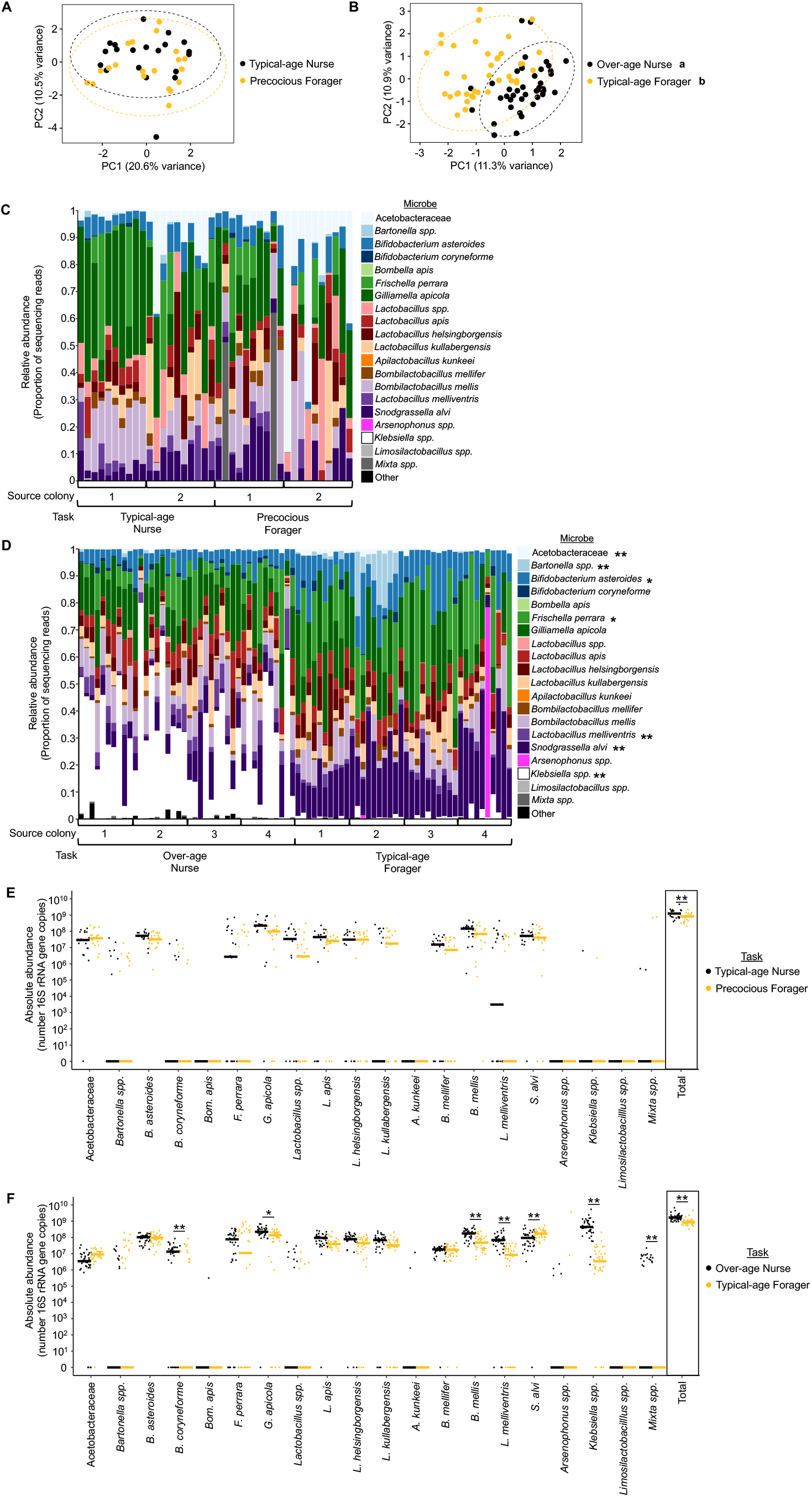
Age-matched nurses and foragers differ in gut microbial community. **(A-B)**Age-matched typical-age nurses and precocious foragers from a single cohort colony did not differ in gut microbial community structure at about one week of age (A), but age-matched over-age nurses and typical-age foragers significantly differed in gut microbial community structure at three weeks of age (B). 1 week: Two-way Permutation MANOVA using Aitchison Distance, Task: F1,39 = 0.92, R^2^ = 0.023, p = 0.519; Source colony: F1,39 = 2.60, R^2^ = 0.064, p = 0.004, Task*Colony: F1,39 = 1.15, R^2^ = 0.028, p = 0.296. n = 10 bees/source colony, 2 source colonies. 3 weeks: Two-way Permutation MANOVA using Aitchison Distance, Task: F1,79 = 6.49, R^2^ = 0.074, p = 0.001; Source colony: F3,79 = 1.82, R^2^ = 0.062, p = 0.001, Task*Colony: F3,79 = 1.23, R^2^ = 0.042, p = 0.084. n = 10 bees/source colony, 4 source colonies. Depicted as PCA plots. Lowercase letters in legends denote statistically significant groups. **(C-D)** Age-matched typical-age nurses and precocious foragers from a single cohort colony did not differ in relative abundance of individual microbial species (C), while age-matched over-age nurses and typical-age foragers differed in relative abundance of five individual microbial species at three weeks of age (D). Depicted as stacked bar plots, with each bar representing a single bee’s microbiome. Asterisks in legend: *, pσ0.05, **, pσ0.01, ANCOM-BC between nurses and foragers. See Supplementary Table 2 for all p-values. **(E-F)** Age-matched typical-age nurses and precocious foragers from a single cohort colony did not differ in absolute abundance of individual microbial species but did differ in the total normalized number of 16S rRNA gene copies (E), while age-matched over-age nurses and typical-age foragers differed in absolute abundance of six microbial species and the total number of 16S rRNA gene copies at three weeks of age (F). 10^x^ number of 16S rRNA gene copies, calculated by multiplying the relative abundance each microbe in each sample (determined through 16S rRNA sequencing) by the normalized number of 16S rRNA gene copies in the sample (determined through qPCR). Depicted as dot plots with all data points plotted, line represents median, n = 10 bees/source colony, 2 source colonies (1 week) or 4 source colonies (3 weeks). *, p≤0.05, **, p≤0.01, Permutation ANOVA Test between nurses and foragers. See Supplementary Table 2 for all p-values.

We next sought to identify individual microbes that differed in abundance between nurse and forager bees to identify candidate microbes specifically associated with division of labor. Since microbes can vary between individuals in relative and/or absolute abundance, we used both of these measures of biological diversity. Both may contribute to ecosystem functioning [103–106], but their relative effects on host behavior are currently unknown [30, 62]. Therefore, we reasoned that using both of these abundance measures would give us a well-rounded sense of individual microbes that are associated with honey bee division of labor. Specifically, we performed a relative abundance analysis (ANCOM-BC [62, 64, 65]) that accounts for the compositional nature of 16S rRNA sequencing data and uses relative abundance measures to estimate the true microbial composition of each gut (hereafter referred to as “relative abundance”). We also combined qPCR with 16S rRNA sequencing data analyses [29, 30] to estimate the absolute abundance of individual microbial taxa in each gut (“absolute abundance”). We found that individual gut microbes significantly differed in both relative and absolute abundance between nurses and foragers, while total microbe abundance did not differ between groups (Fig. 1C-D, Supplementary Table 1, Table 1). These results indicate that nurse and forager bees differ significantly in their gut microbial communities. They also identify candidate microbes that may be associated with division of labor.

**Table 1:** Summary table of individual microbe abundance results. Microbes that significantly differed in relative (left) and absolute (right) abundance measures in nurses relative to foragers in typical colonies and individual single cohort colony replicates. Higher abundance, higher in abundance in nurses relative to foragers (pink). Lower abundance, lower in abundance in nurses relative to foragers (blue)

| Microbe | Relative abundance |  |  | Absolute abundance |  |  |
| --- | --- | --- | --- | --- | --- | --- |
|  | Typical colonies | SCC Replicate 1 (week 3) | SCC Replicate 2 (week 3) | Typical colonies | SCC Replicate 1 (week 3) | SCC Replicate 2 (week 3) |
| Acetobacteraceae |  | lower abundance | lower abundance | lower abundance |  | lower abundance |
| *Bartonella* spp. |  | lower abundance |  |  |  |  |
| *B. asteroides* |  | lower abundance | lower abundance |  |  |  |
| *B. coryneforme* |  |  |  |  | higher abundance |  |
| *Bom. apis* |  |  |  |  |  |  |
| *F. perrara* |  | lower abundance |  |  |  |  |
| *G. apicola* |  |  |  |  | higher abundance | higher abundance |
| *Lactobacillus* spp. |  |  |  | higher abundance |  |  |
| *L. apis* |  |  |  |  |  | higher abundance |
| *L. helsingborgensis* | higher abundance |  |  |  |  |  |
| *L. kullabergensis* |  |  |  | higher abundance |  | higher abundance |
| *A. kunkeei* |  |  |  |  |  |  |
| *B. mellifer* |  |  |  |  |  |  |
| *B. mellis* | higher abundance |  |  |  | higher abundance | higher abundance |
| *L. melliventris* | higher abundance | higher abundance | higher abundance | higher abundance | higher abundance |  |
| *S. alvi* |  | lower abundance |  | lower abundance | lower abundance |  |
| *Arsenophonus* spp. |  |  |  |  |  |  |
| *Klebsiella* spp. |  | higher abundance | higher abundance | higher abundance | higher abundance | higher abundance |
| *Limosilactobacillus* spp. |  |  |  |  |  |  |
| *Mixta* spp. |  |  |  |  | higher abundance |  |

**Table 2:**
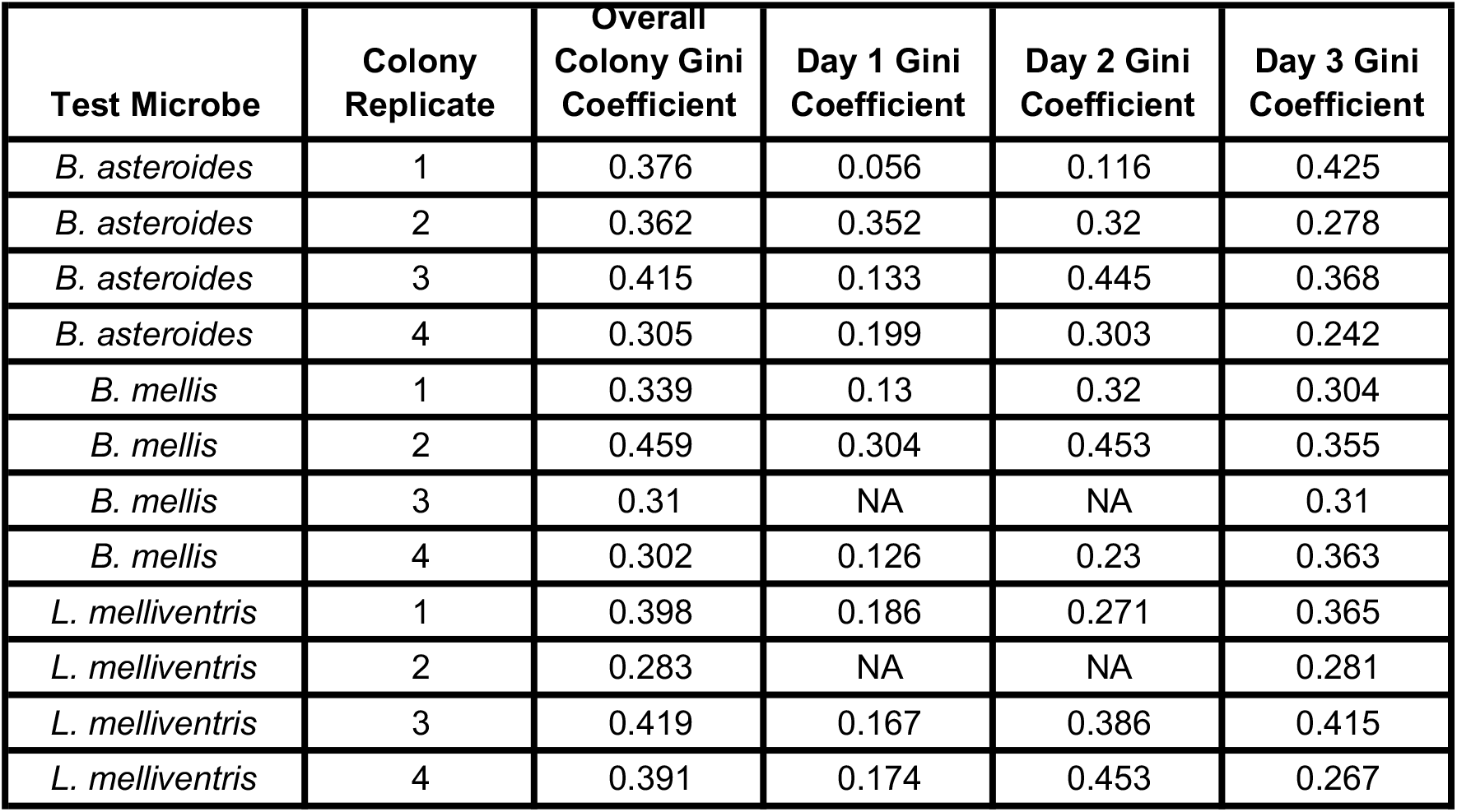
Gini coefficients for each experimental colony depicted in. **Figures 4-6**. Gini coefficients, representing the degree of skew in foraging intensity between individual foraging bees, where 0 represents complete equality in foraging intensity between individuals and 1 represents complete inequality in foraging intensity between individuals, measured for each experimental colony across all days (“Overall Colony Gini Coefficient”), and for each experimental colony on each individual day (“Day X Gini Coefficient”). NAs represent days with bad weather in which foraging did not occur.

### Differences in gut microbial community composition between nurses and foragers are dependent upon differences in behavioral state

Under typical colony conditions, nurses and foragers differ in chronological age. However, honey bee division of labor is flexible and responsive to changing colony needs. For example, if a colony experiences a shortage of older bees, some younger individuals will begin to forage at an early age, while a shortage of younger bees will cause bees to continue to act as nurse bees despite advancing chronological age [35–39]. To determine whether the gut microbial differences between nurses and foragers depend on differences in behavioral state and/or age, we exploited this adaptive plasticity to create SCCs, a well-established experimental approach that separates these two factors [36, 37, 51]. In two SCC replicates, we compared gut microbiomes between age-matched nurse and forager bees when they were about one week of age (representing typical-age nurses and precocious foragers) and three weeks of age (representing over-age nurses and typical-age foragers).

Age-matched typical-age nurses and precocious foragers did not significantly differ in gut microbial community structure (Fig. 2A, Supplementary Fig. 4A), while age-matched over-age nurses and typical-age foragers did (Fig. 2B, Supplementary Fig. 4B). Similar results were obtained for the abundance of individual microbial taxa: no microbes consistently differed in abundance between typical-age nurses and precocious foragers across the two SCC replicates (Fig. 2C, Fig. 2E, Supplementary Fig. 4C, Supplementary Fig. 4E, Supplementary Table 2, Supplementary Table 3), while four microbes consistently differed in relative abundance and three microbes consistently differed in absolute abundance between over-age nurses and typical-age foragers across the two SCC replicates (Fig. 2D, Fig. 2F, Supplementary Fig. 4D, Supplementary Fig. 4F, Supplementary Table 2, Supplementary Table 3). In addition, the total absolute abundance of microbes differed between age-matched typical-age nurses and precocious foragers in one colony replicate (Fig. 2E, Supplementary Fig. 4E), and between age-matched over-age nurses and typical-age foragers (Fig. 2F, Supplementary Fig. 4F). Overall, these findings indicate that age-dependent differences in behavioral state are associated with differences in gut microbial communities between nurses and foragers.

Together, results from the typical colony and three-week old SCC indicate that of the honey bee-associated microbes, *Bombilactobacillus mellis* (previously *Lactobacillus mellis*) and *Lactobacillus melliventris* were consistently (i.e. identified in three or more analyses across the two abundance measures) more abundant in nurses relative to foragers (Table 1), while Acetobacteraceae and *Snodgrassella alvi* were consistently less abundant in nurses relative to foragers (Table 1). In SCCs alone, *Bifidobacterium asteroides* showed consistent lower relative abundance in over-age nurses compared to typical-age foragers (Table 1) and *Gilliamella apicola* (Table 1) showed consistent higher absolute abundance in over-age nurses compared to typical-age foragers. Such robust associations highlight these six microbes as candidate microbes whose abundances may have causal effects on division of labor in honey bees. In addition, *Klebsiella spp.*, a taxonomic group not typically considered a honey bee-associate, was more abundant in over-age nurses compared to typical-age foragers (Fig. 1-2, Supplementary Fig. 4, Table 1). This microbial species group has previously been found to be prevalent in nurse bees and, as an environmentally derived potential pathogen to bees, was likely picked up outside of the hive by foragers and disseminated to nurses within the hive [30]. As previously suggested, it may accumulate in the guts of nurses, which are typically heavier and support higher bacterial loads than forager guts [30]. This possibility is consistent with our finding that *Klebsiella* was absent in typical-age nurses and precocious foragers but present in over-age nurses and typical-age foragers, likely indicating that *Klebsiella* was inadvertently introduced to the colony after one week, at which time it began accumulating in the guts of over-age nurses, who have higher bacterial loads than typical-age foragers (Fig. 2, Supplementary Fig. 4).

The lack of gut microbial community differences between one-week old SCC nurses and foragers suggests that differences in behavioral maturation rate are not associated with differences in gut microbial community composition. Rather, the observed microbiome differences between over-age nurses and typical-age foragers suggest that associations between gut microbiome and behavioral state depend on worker experience, possibly, as others have suggested, due to task-related differences in environment, diet, and/or metabolic needs [30–32]. Since SCCs are composed of bees of a single age cohort, three-week old SCCs are composed of bees that have been performing their respective tasks for prolonged periods (1-2 weeks longer) compared to one-week old SCCs. Therefore, we hypothesized that task experience plays an important role in gut microbial community composition. In the next section, we tested this by manipulating foraging experience since it is known to have strong effects on behavior, brain chemistry, brain structure, and brain gene expression [75, 97, 107, 108].

### Foraging experience influences gut microbial community structure, while behavioral state influences the abundance of most individual microbes

In order to determine the causal effects of behavioral state and foraging experience on honey bee gut microbiome composition, we controlled for behavioral state while manipulating foraging experience. To do this, we created “big-back colonies” in which some foragers were freely able to leave the colony (“active foragers”), while other bees, who appeared at the entrance and showed an inclination to forage, were prevented from ever leaving the hive due to the presence of a thick plastic tag on their backs (“inactive foragers”) [51, 53]. We were thus able to compare bees of the same age and behavioral state that differed in foraging experience.

We found that nurses and inactive foragers did not differ in gut microbial community structure, while active foragers and inactive foragers did (Fig. 3A). This finding indicates that foraging experience plays an important role in defining differences in gut microbial community structure.

**Figure 3.**
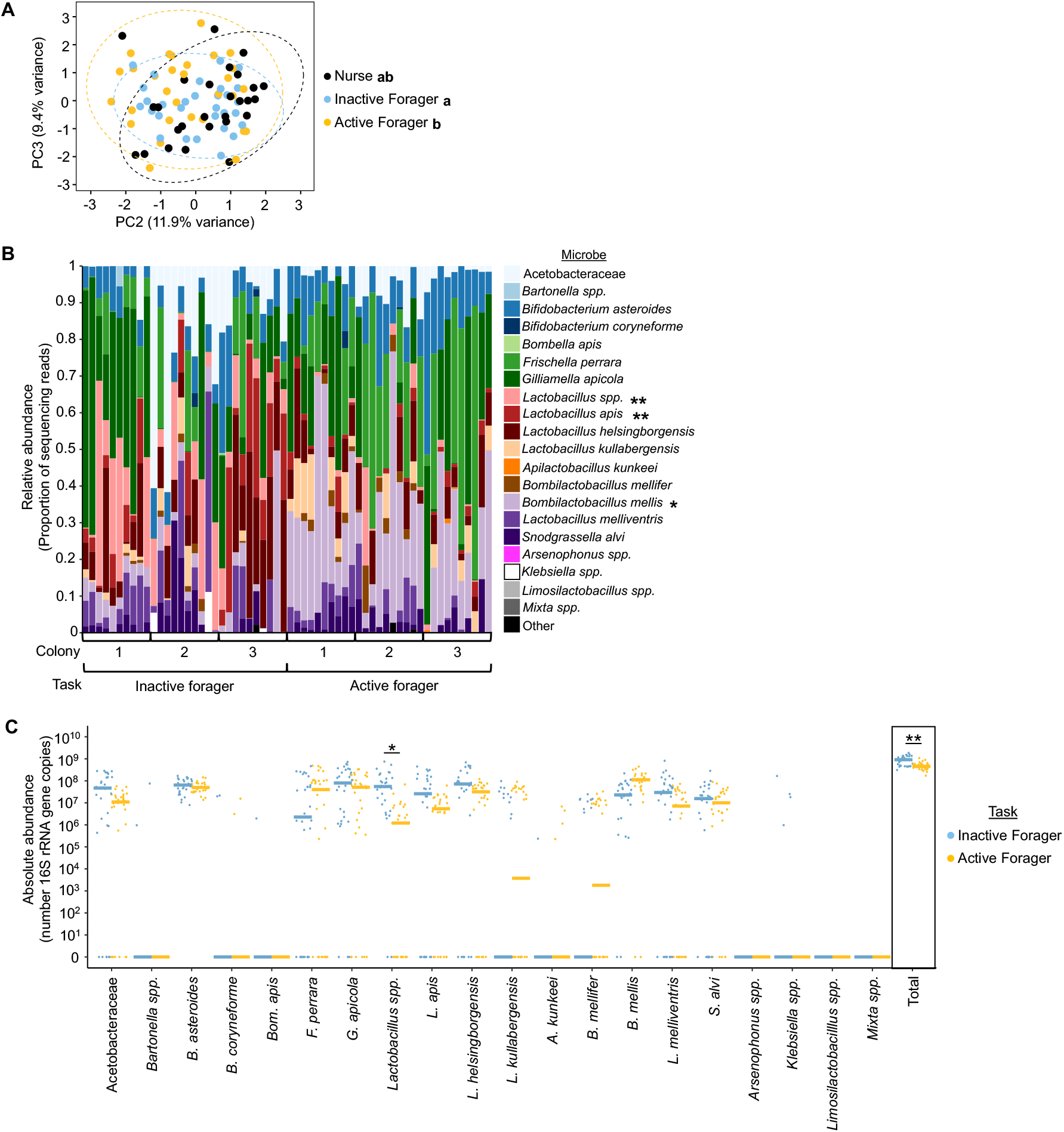
Age-matched inactive and active foragers differ in gut microbial community structure, but not the abundance of individual microbes. **(A)**Age-matched nurses and inactive foragers were similar in gut microbial community structure, while age-matched active and inactive foragers differed in gut microbial community structure. Two-way Permutation MANOVA using Aitchison Distance, Task: F2,86 = 1.84, R^2^ = 0.038, p = 0.011; Colony: F2,86 = 6.02, R^2^ = 0.124, p = 0.001, Task*Colony: F4,86 = 0.798, R^2^ = 0.033, p = 0.883. n = 10 bees/colony, 3 colonies. Depicted as PCA plot. Lowercase letters in legends denote statistically significant groups as determined by Pairwise Permutation MANOVA. **(B)** Age-matched inactive and active foragers did not differ in the relative abundance of individual microbial species (B). Depicted as stacked bar plots, with each bar representing a single bee’s microbiome. Asterisks in legend: *, pσ0.05, **, pσ0.01, ANCOM-BC between inactive and active foragers. See Supplementary Table 4 for all p-values. **(C)** Age-matched inactive and active foragers did not differ in absolute abundance of individual microbial species but did differ in the total normalized number of 16S rRNA gene copies. 10^x^ number of 16S rRNA gene copies, calculated by multiplying the relative abundance each microbe in each sample (determined through 16S rRNA sequencing) by the normalized number of 16S rRNA gene copies in the sample (determined through qPCR). Depicted as dot plots with all data points plotted, line represents median, n = 10 bees/colony, 3 colonies. *, pσ0.05, **, pσ0.01, Permutation ANOVA Test between inactive and active foragers. See Supplementary Table 4 for all p-values.

**Figure 4.**
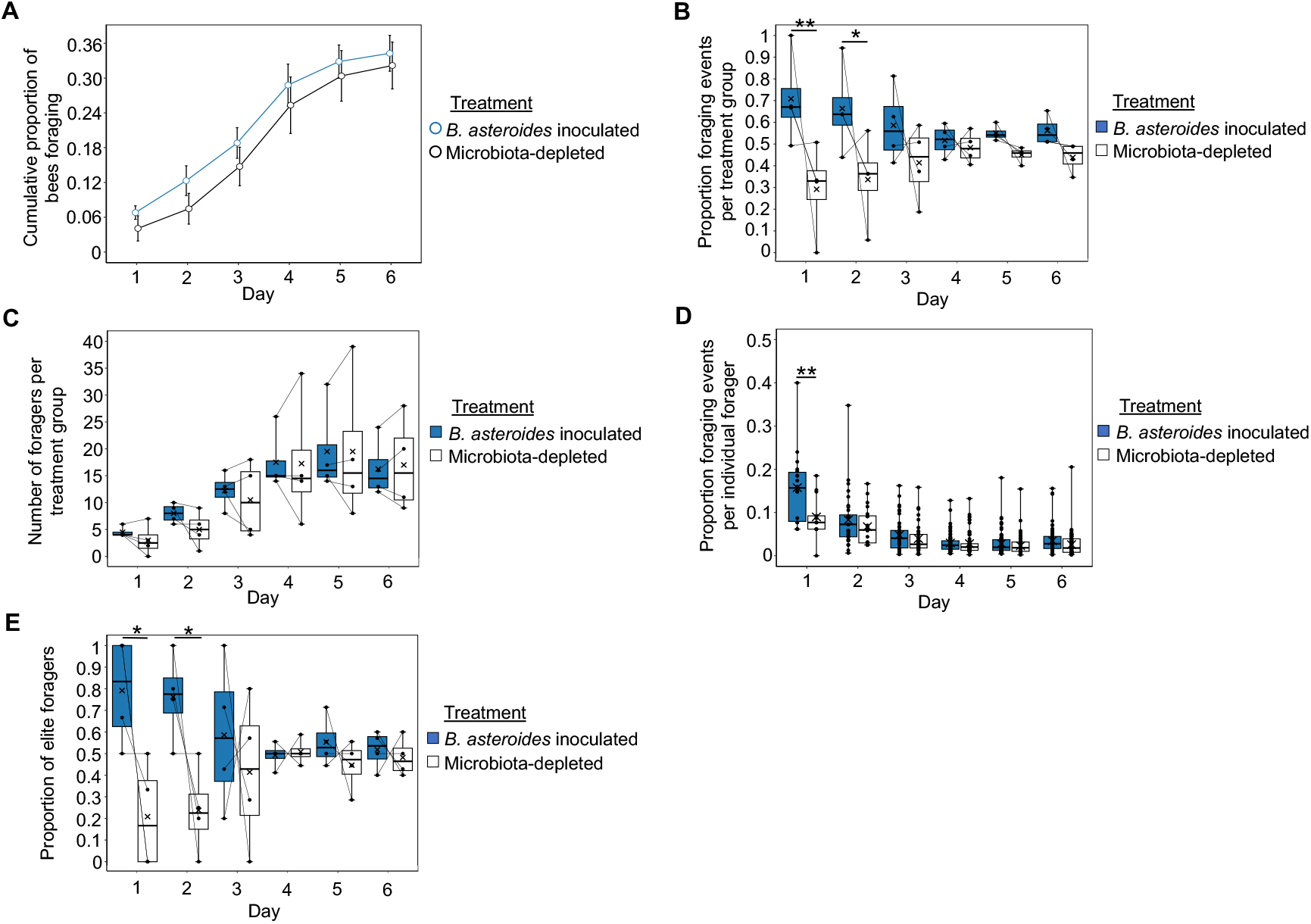
Bees inoculated with *B. asteroides* do not differ from microbiota-depleted bees in behavioral maturation (age at onset of foraging), but do differ in foraging intensity. **(A)***B. asteroides* inoculated bees did not differ from microbiota-depleted bees in age at onset of foraging. Cox Proportional Hazards, z = -0.331, p = 0.741. **(B)** *B. asteroides* inoculated foragers, as a group, performed the majority of foraging trips for the colony on the first and second days after experimental colony formation. Linear Mixed Effects Model, Treatment: F1,33 = 21.16, p < 0.001, Day: F5,33 = 0, p = 1, Treatment*Day: F5,33 = 1.987, p = 0.107. See Supplementary Table 5 for Pairwise comparisons. **(C)** *B. asteroides* inoculated bees represented a similar number of foragers as microbiota-depleted bees on all days after experimental colony formation. Linear Mixed Effects Model, Treatment: F1,33 = 0.409, p = 0.527, Day: F5,33 = 12.095, p < 0.001, Treatment*Day: F5,33 = 0.140, p = 0.982. See Supplementary Table 5 for Pairwise comparisons. Individual *B. asteroides* inoculated foragers performed a majority of foraging trips for the colony the first day after experimental colony formation. Generalized Linear Mixed Effects Model with log-normal distribution, Treatment: ξ^2^ = 4.985, p = 0.026, Day: ξ^2^ = 382.933, p < 0.001, Treatment*Day: ξ^2^ = 26.315, p < 0.001. See Supplementary Table 5 for Pairwise comparisons. **(D)** *B. asteroides* inoculated bees represented a higher proportion of elite foragers than microbiota-depleted bees on the first and second days after experimental colony formation. Linear Mixed Effects Model, Treatment: F1,33 = 15.799, p < 0.001, Day: F5,33 = 0, p = 1, Treatment*Day: F5,33 = 3.151, p = 0.020. See Supplementary Table 5 for Pairwise comparisons. (A) depicted as survival plot. All other data depicted as box plots with data points plotted, thick horizonal line represents median, x represents mean, whiskers represent the minimum and maximum values, thin lines connect paired points for each colony, n = 4 colonies. Asterisks used to denote comparisons between treatment groups on each day only: *, pσ:0.05, **, pσ:0.001.

**Figure 5.**
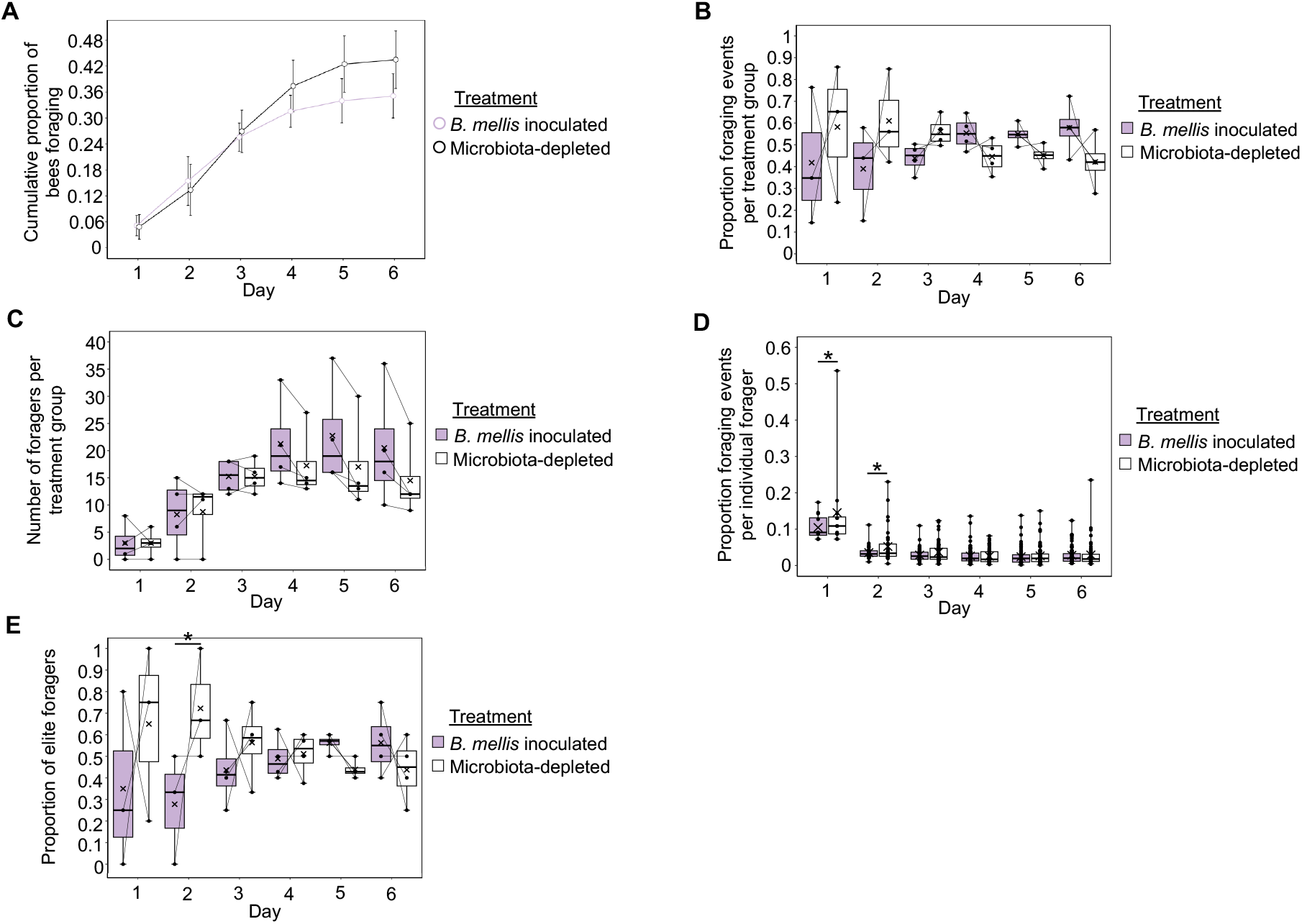
Bees inoculated with *B. mellis* do not differ from microbiota-depleted bees in behavioral maturation (age at onset of foraging), but do differ in foraging intensity. **(A)***B. mellis* inoculated bees did not differ from microbiota-depleted bees in age at onset of foraging. Cox Proportional Hazards, z = 1.525, p = 0.127. **(B)** *B. mellis* inoculated foragers, as a group, performed a similar proportion of foraging trips for the colony as microbiota-depleted bees all days after experimental colony formation, with a marginal effect of performing a minority of foraging trips for the colony on the second day after colony formation. Linear Mixed Effects Model, Treatment: F1,29.091 = 0.036, p = 0.851, Day: F5,29.719 = 0, p = 1, Treatment*Day: F5,29.091 = 1.981, p = 0.111. See Supplementary Table 5 for Pairwise comparisons. **(C)** *B. mellis* inoculated bees represented a similar number of foragers as microbiota-depleted bees on all days after experimental colony formation. Linear Mixed Effects Model, Treatment: F1,33 = 3.183, p = 0.084, Day: F5,33 = 14.918, p < 0.001, Treatment*Day: F5,33 = 0.765, p = 0.582. See Supplementary Table 5 for Pairwise comparisons. **(D)** Individual *B. mellis* inoculated foragers performed a minority of foraging trips for the colony on the first and second days after experimental colony formation. Generalized Linear Mixed Effects Model with log-normal distribution, Treatment: ξ^2^ = 3.626, p = 0.057, Day: ξ^2^ = 255.497, p < 0.001, Treatment*Day: ξ^2^ = 6.552, p = 0.256, See Supplementary Table 5 for Pairwise comparisons. No statistical outliers were detected in these data. **(E)** *B. mellis* inoculated bees represented a lower proportion of elite foragers than microbiota-depleted bees on the second day after experimental colony formation. Linear Mixed Effects Model, Treatment: F1,29.091 = 1.896, p = 0.179, Day: F5,29.719 = 0, p = 1, Treatment*Day: F5,29.091 = 2.183, p = 0.083. See Supplementary Table 5 for Pairwise comparisons. (A) depicted as survival plot. All other data depicted as box plots with data points plotted, thick horizonal line represents median, x represents mean, whiskers represent the minimum and maximum values, thin lines connect paired points for each colony, n = 4 colonies. Asterisks used to denote comparisons between treatment groups on each day only: *, pσ:0.05, **, pσ:0.001.

**Figure 6.**
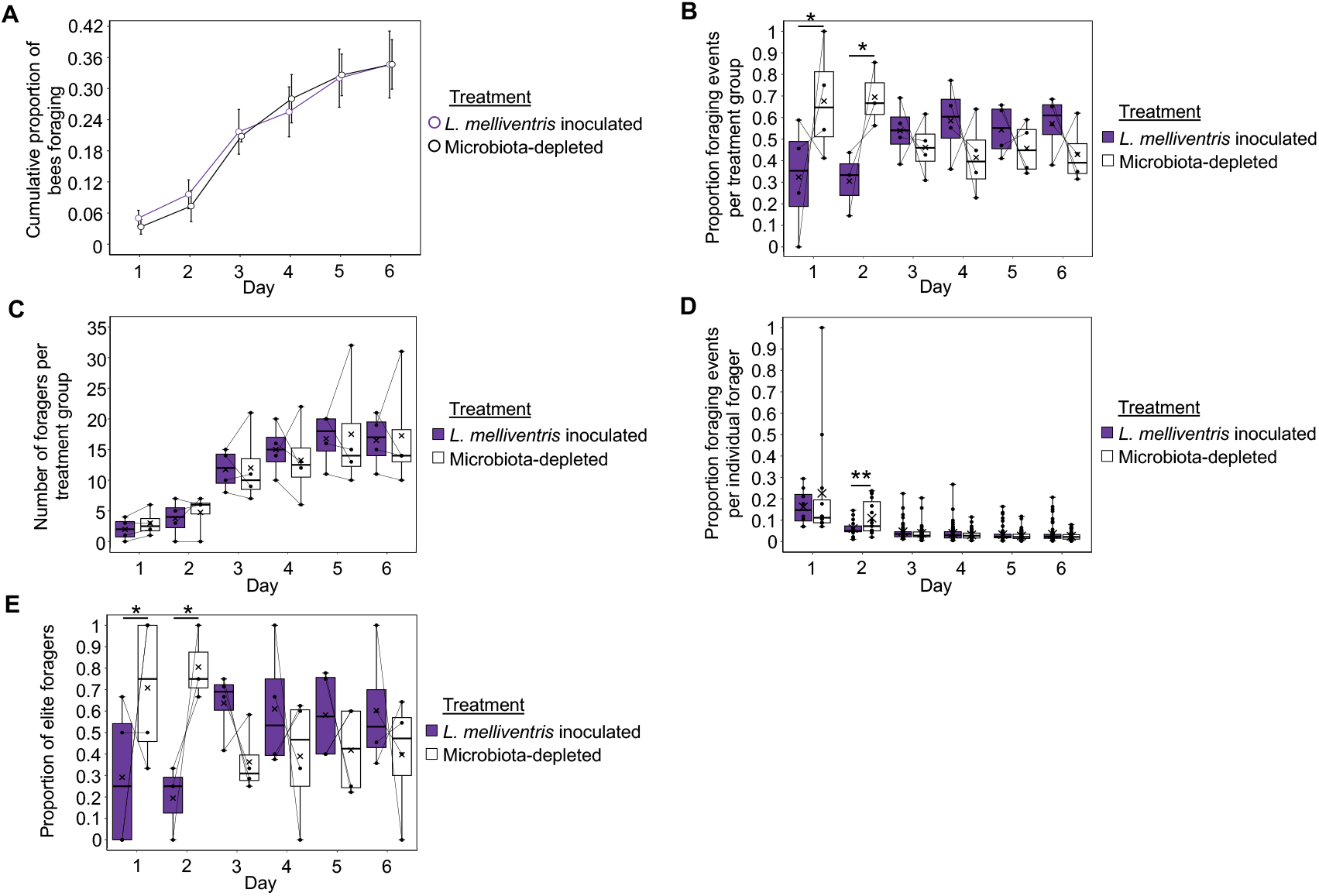
Bees inoculated with *L. melliventris* do not differ from microbiota-depleted bees in behavioral maturation (age at onset of foraging), but do differ in foraging intensity. **(A)***L. melliventris* inoculated bees did not differ from microbiota-depleted bees in age at onset of foraging. Cox Proportional Hazards, z = -0.435, p = 0.664. **(B)** *L. melliventris* inoculated foragers, as a group, performed the minority of foraging trips for the colony on the first and second days after experimental colony formation. Linear Mixed Effects Model, Treatment: F1,31.021 = 0.351, p = 0.558, Day: F5,31.443 = 0, p = 1, Treatment*Day: F5,31.021 = 4.169, p = 0.005. See Supplementary Table 5 for Pairwise comparisons. **(C)** *L. melliventris* inoculated bees represented a similar number of foragers as microbiota-depleted bees on all days after experimental colony formation. Linear Mixed Effects Model, Treatment: F1,33 = 0.079, p = 0.781, Day: F5,33 = 18.923, p < 0.001, Treatment*Day: F5,33 = 0.132, p = 0.984. See Supplementary Table 5 for Pairwise comparisons. **(D)** Individual *L. melliventris* inoculated foragers performed a minority of foraging trips for the colony the second day after experimental colony formation. Generalized Linear Mixed Effects Model with log-normal distribution, Treatment: ξ^2^ = 0.141, p = 0.707, Day: ξ^2^ = 334.831, p < 0.001, Treatment*Day: ξ^2^ = 18.336, p = 0.003, See Supplementary Table 5 for Pairwise comparisons. No statistical outliers were detected in these data. **(E)** *L. melliventris* inoculated bees represented a lower proportion of elite foragers than microbiota-depleted bees on the first and second days after experimental colony formation. Linear Mixed Effects Model, Treatment: F1,31.021 = 0.0006, p = 0.981, Day: F5,31.443 = 0, p = 1, Treatment*Day: F5,31.021 = 3.961, p = 0.007. See Supplementary Table 5 for Pairwise comparisons. (A) depicted as survival plot. All other data depicted as box plots with data points plotted, thick horizonal line represents median, x represents mean, whiskers represent the minimum and maximum values, thin lines connect paired points for each colony, n = 4 colonies. Asterisks used to denote comparisons between treatment groups on each day only: *, p≤0.05, **, p≤0.001.

When we compared the abundance of individual taxa between age-matched active and inactive foragers, we found that unclassified *Lactobacillus spp.* were higher in relative and absolute abundance in inactive foragers compared to active foragers, *L. apis* was higher in relative abundance in inactive foragers compared to active foragers, and *B. mellis* was lower in relative abundance in inactive foragers compared to active foragers (Fig. 3B-C, Supplementary Table 4). In addition, inactive foragers had a higher absolute abundance of total microbes compared to active foragers (Fig. 3C). This effect seems to be driven by a low absolute total abundance in the active foragers, as the inactive foragers had a median absolute total abundance similar to foragers in typical colonies and three-week old SCCs (Supplementary Tables 1-4). Therefore, while inactive foragers are unable to leave the colony to defecate, differences in microbiome between inactive and active foragers are unlikely to be due to an unhealthy accumulation of bacterial cells in inactive forager guts.

The observed difference in *Lactobacillus spp.* abundance between inactive and active foragers matches our earlier finding that this taxonomic group decreases in absolute abundance between nurses and foragers in typical colonies (Fig. 1D), indicating that this microbe group may be associated with task experience. Interestingly, in our earlier analyses *B. mellis* was higher in relative and absolute abundance measures in nurses compared to foragers (Fig. 1, Fig. 2, Supplementary Fig. 4, Table 1). While we did not find an effect on absolute abundance here, it is surprising that *B. mellis* is higher in relative abundance in active foragers compared to inactive foragers. While all of our analyses support a connection between *B. mellis* and division of labor, this relationship may be more complex than originally considered from nurse/forager results alone. The other division of labor associated microbes (Table 1) did not differ in abundance between inactive and active foragers.

Overall, these results suggest that the abundance of some microbes is determined by the experience of foraging outside of the hive, while the abundance of most individual microbes is defined by behavioral state. Together with our SCC results, which indicate that differences in the abundance of individual gut microbes may be dependent upon worker experience (Fig. 2D,F, Supplementary Fig. 4D,F), we speculate that honey bee gut microbes and foraging behaviors interact through a feedback model. For example, gut microbes may change in abundance due to foraging experience, possibly incrementally (and thus indiscernibly) at first and building with more experience, and this change in microbiome may then feed back to support host foraging behaviors [32]. In the next section, we test the latter part of this model by assessing the effects of inoculation with individual microbes on host foraging behavior.

### Inoculation with individual microbes affects group- and individual-level foraging intensity

To determine if gut microbes play a causal role in honey bee division of labor, we performed single microbe inoculations with three out of the six individual honey bee microbes robustly associated with division of labor as described above: *B. asteroides*, *B. mellis* and *L. melliventris* (Supplementary Fig. 5A-C). In addition to the strong patterns of association reported above, we chose these three microbes due to *B. asteroides*’ known effects on host physiology with links to behavior, the high metabolic output associated with taxa in Lactobacillaceae [46], since gut microbial metabolites likely contribute to the microbiota-gut-brain axis [7], and due to these three species’ previously published associations with honey bee behavioral task [30–32, 109]. Therefore, we reasoned that these three species represented strong candidates to test for causal effects on honey bee division of labor. Using an automated behavioral tracking system [71], we determined the effect of inoculation with these three microbes on two measures of foraging behavior important in division of labor in honey bees: rate of behavioral maturation (age at onset of foraging) and foraging intensity (amount of foraging trips performed by a group or individual) [44]. Our results indicate effects only during the first two days of behavioral monitoring, for reasons that we speculate about in the Discussion.

*A. asteroides* inoculated bees did not differ from microbiota-depleted bees in behavioral maturation rate (Fig. 4A) across experimental replicates, but did differ in foraging intensity. Overall, *B. asteroides* inoculated bees performed the majority of the foraging trips for the colony on the first and second days after colony formation (Fig. 4B, Supplementary Table 5). These group-level effects were not due to a difference in number of foragers between inoculated and microbiota-depleted bees (Fig. 4C, Supplementary Table 5). Rather they were at least partly due to a difference in individual-level foraging intensity, as individual *B. asteroides* inoculated foragers performed a majority of the foraging trips for the colony on the first day after colony formation (Fig. 4D, Supplementary Table 5). Likewise, *B. asteroides* experimental colonies displayed a degree of skew in foraging intensity among individuals (Table 2) and *B. asteroides* inoculated bees represented a higher proportion of “elite forager” bees, i.e., a small subset of foragers that performed ≥50% of the colony’s foraging trips [43, 75], relative to microbiota-depleted bees on the first and second days after colony formation (Fig. 4E, Supplementary Table 5).

Similar to *B. asteroides* inoculated bees, *B. mellis* inoculated bees did not differ from microbiota-depleted bees in behavioral maturation rate (Fig. 5A). By contrast, *B. mellis* inoculation did not have an effect on group-level foraging intensity (Fig. 5B, Supplementary Table 5) or number of foragers between inoculated and microbiota-depleted bees (Fig. 5C, Supplementary Table 5). Rather, *B. mellis* inoculation influenced individual-level foraging intensity, as *B. mellis* inoculated bees performed a minority of foraging trips for the colony on the first and second days after colony formation (Fig. 5D, Supplementary Table 5). Additionally, *B. mellis* inoculation caused a skew in foraging intensity between individual foragers (Table 2), as *B. mellis* inoculated bees represented a lower proportion of elite foragers than microbiota-depleted bees on the second day after colony formation (Fig. 5E, Supplementary Table 5).

Similar to *B. asteroides* and *B. mellis* inoculated bees, *L. melliventris* inoculated bees did not differ from microbiota-depleted bees in behavioral maturation rate (Fig. 6A). However, *L. melliventris* inoculated bees had an opposite effect on foraging intensity from *B. asteroides* inoculated bees: *L. melliventris* inoculated bees performed the minority of foraging trips for the colony on the first and second days after colony formation (Fig. 6B, Supplementary Table 5). This group-level effect was not due to a difference in number of foragers between inoculated and microbiota-depleted bees (Fig. 6C, Supplementary Table 5). Rather, it was at least partly due to a difference in individual-level foraging intensity, as individual *L. melliventris* inoculated foragers performed a minority of the foraging trips for the colony on the second day after colony formation (Fig. 6D, Supplementary Table 5). Likewise, *L. melliventris* colonies displayed a degree of skew in foraging intensity between individual foragers (Table 2), and *L. melliventris* bees represented a lower proportion of elite foragers than microbiota-depleted bees on the second day after colony formation (Fig. 6E, Supplementary Table 5).

Overall, these results indicate that inoculation with individual gut microbes is sufficient to cause increases or decreases in foraging intensity between worker bees at both the group- and individual-level.

## Discussion

Here, we show that the gut microbiome plays an important role in modulating foraging behavior in the honey bee. Specifically, our results indicate that differences in honey bee gut microbial communities influence honey bee division of labor by affecting foraging intensity, but not rate of behavioral maturation. This conclusion is supported by both microbe abundance and behavioral results: *B. asteroides* is found in higher relative abundance in foragers compared to nurses, and inoculation with this species caused bees to have increased foraging intensity compared to microbiota-depleted controls. Similarly, *B. mellis* and *L. melliventris* are found lower in abundance in foragers compared to nurses, and inoculation with either of these species caused bees to have decreased foraging intensity compared to microbiota-depleted controls.

From a technical perspective, the observation that single microbe inoculations can cause either an increase or a decrease in foraging intensity indicates that it is unlikely that the results reported here are due to an artifact of the microbiological or behavioral methods we used. In addition, the congruence of our microbe abundance and behavioral results indicate that single microbe inoculations, although they do not represent natural microbiome variations, provide a useful method for functional analysis.

Overall, our behavioral results show the strongest effects on the first and second days after experimental colony formation, indicating that the most robust effects of microbiome on behavior occur within the first few days after inoculation. We suspect that this is due to changing social dynamics in the experimental honey bee colonies over the six day behavioral tracking period. Although we did not measure gut microbiomes post colony introduction, it is likely that changes in the gut microbiomes of the experimental bees occurred after colony formation due to “homogenization” between individuals due to “trophallaxis,” the sharing of gut fluids that contain nutritive and signaling molecules, and/or colonization by microbes acquired from the colony environment or non-experimental bees. Therefore, experimental treatment effects on behavior may be weak or non-existent on later experimental days. Changes in colony dynamics, such as the pool of available foragers, may also account for weak behavioral effects observed later in the experiment. Foraging is an energetically costly task and is associated with high mortality compared to in-hive behaviors [110–112]. Indeed, across all types of experimental colonies in our studies, foragers that began foraging early after colony formation had an earlier last day of foraging (which likely corresponds with day of death [44, 74]) (Supplementary Fig. 5D-F). This indicates that our experimental colonies had a typical high forager turnover.

We observed additional interesting changes in colony foraging force due to foraging activity. In the case of *B. mellis* experimental colonies, only microbiota-depleted foragers that began foraging on the first day had a significantly earlier last day of foraging compared to later foragers (Supplementary Fig. 5E). In addition, microbiota-depleted elite foragers had a significantly earlier first day of foraging compared to *B. mellis* inoculated elite foragers (Supplementary Fig. 5G). These results indicate that in *B. mellis* experimental colonies, forager turnover was associated with treatment such that microbiota-depleted foragers had their foraging career at younger ages compared to *B. mellis* inoculated foragers. A similar but non-significant trend in elite forager onset was also observed in *L. melliventris* colonies (Supplementary Fig. 5G). We suggest that this likely accounts for the (non-significant) switch in group-level and elite foraging intensity seen on the later experimental days in *B. mellis* (Fig. 5B, Fig. 5E) and *L. melliventris* (Fig. 6B, Fig. 6E), but not *B. asteroides* (Fig. 4) experimental colonies.

Previous studies reported correlational differences in gut microbiome associated with honey bee behavioral task [30–32] and behavioral maturation [50]. These studies also reported associations of *Bifidobacterium*, *B. mellis* and *L. melliventris* with behavioral task [30, 32] and ontogeny [31], supporting a link between these microbes and division of labor in honey bees. Here we found that *B. mellis* and *L. melliventris* were higher in abundance in nurses compared to foragers, matching results from [30, 32]. However, while we found that *B. asteroides* was higher in relative abundance in typical-age foragers relative to over-age nurses, results in [30, 32] found that *Bifidobacterium* was higher in relative [32] and absolute abundance [30] in nurses compared to foragers. This difference may be due to the difference in taxonomic level used in analysis between these studies and ours, as we found that other *Bifidobacterium* species were higher in abundance in nurses compared to foragers (trend in Fig. 1C-D and Fig. 2D, significant in Fig. 2F). In addition, these studies found an increase in *S. alvi* with age [31], and a higher abundance of *Gilliamella* and *Klebsiella* associated with nurses [30], which match our abundance results (Table 1). Here, we provide evidence that at least some of these gut microbes play a causal role in defining behavioral differences between honey bees. Therefore, the congruence of our abundance results with other studies indicate that our behavioral results are likely generalizable across populations of honey bees.

Our findings are consistent with a previously proposed model in which positive feedback interactions cause adaptive changes in both gut microbiome composition and host behavior [32]. Under this model, we speculate that gut microbes could change in abundance due to time spent foraging, likely in association with environmental exposure and/or changes in diet and metabolic needs. According to this speculation, the change in microbiome would then feed back to influence host foraging behaviors. We suggest that changes in gut microbe abundance (e.g. increases in *B. asteroides* with decreases in *B. mellis* and *L. melliventris*) and host behavior interact to increase foraging intensity. Modeling interactions in this way can provide the basis for future studies that probe the neural and metabolic mechanisms by which *B. asteroides*, *B. mellis* and *L. melliventris* affect host behavior.

Previous studies have shown that gut microbes may influence host behavior through a variety of mechanisms, including production of metabolites that cause changes in host brain gene expression or host production of neurotransmitters and hormones [7]. Within honey bees, *Bifidobacterium*, *Bombilactobacillus* Firm-4 and *Lactobacillus* Firm-5 are the major fermenters of pollen-derived compounds and have large effects on the abundance of metabolites in the hindgut [46]. In addition, *B. asteroides* inoculation has been associated with an increased production of two juvenile hormone derivatives [46]. Changes in gene expression and the production of neurotransmitters and hormones, most notably juvenile hormone, are associated with the regulation of honey bee division of labor [42, 44, 74, 95–98, 101, 107, 113, 114], and therefore it is possible that gut microbes influence honey bee division of labor through these pathways. Likewise, previous studies indicate that some host factors associated with honey bee division of labor, such as social interactions and diet, may also modulate gut microbiome composition, including the abundance of these three microbes [7, 30, 48, 115, 116]. Future research will work to identify specific mechanisms by which honey bees and their gut microbiomes interact to influence worker behavior. Overall, studies using the honey bee as a model have the potential to further elucidate the microbiota-gut-brain axis, and achieve a more comprehensive understanding of interkingdom interactions.

Together, our results suggest that in naturally occurring bee populations, increases in *B. asteroides* with simultaneous decreases in *B. mellis* and *L. melliventris* may act synergistically to increase the intensity of foraging behavior in individual bees. Honey bees, with a complex eusocial life history, live in large perennial colonies and therefore collect large amounts of nectar and pollen to ensure colony survival during periods when floral resources are not available. Thus, the influence of these three microbes on honey bee foraging intensity indicates that host-microbe interactions likely help sustain the perennial lifestyle of this eusocial insect. Future studies may address the interaction between these three microbes, other members of the honey bee microbiome, and between the microbiome and the host in order to gain a more naturalistic and comprehensive understanding of the role of the gut microbiome in social insect division of labor.

## Acknowledgements

We thank Nathan Beach, Sarah Magdalena Murphree, Sarai Stuart, Laura Kilikevicius, Rashmi Bajaj, Maha Syed and Anthony Cantu for field assistance, Sue Cobey and Dr. Osman Kaftanoglu for SDI queen rearing and inseminations, Ahmed Burki for assistance making qPCR standard plasmids, Alvaro Hernandez, Chris Wright and staff at the Carver Biotechnology Center for sequencing services, members of the Computer Network Resource Group of the Carl R. Woese Institute for Genomic Biology (UIUC) for computational support, Konrad Wilk for annotating missed barcodes, Richard B. Devine for contributions to video annotation software development, and the workers of Turker Nation for annotating honey bee trajectories. Funding: US National Science Foundation Division of Integrative Organismal Systems award number 2120378, PI: GER.

## Data Availability Statement

16S rRNA sequencing data are available from the NCBI Sequence Read Archive (PRJNA958253) and all other data are available from Mendeley Data (DOI: 10.17632/f2s47y3nhn). Computer code for the barcode detector is available at https://github.com/gernat/btools.

## References

1. Zilber-Rosenberg I, Rosenberg E. Role of microorganisms in the evolution of animals and plants: the hologenome theory of evolution. FEMS Microbiol Rev 2008; 32: 723–735.

2. Cryan JF, Dinan TG. Mind-altering microorganisms: the impact of the gut microbiota on brain and behaviour. Nat Rev Neurosci 2012; 13: 701–712.

3. Flint HJ, Karen P, Louis P, Duncan SH. The role of the gut microbiota in nutrition and health. Nat Rev Gastroenterol Hepatol 2012; 9: 577–589.

4. Hooper L V., Littman DR, Macpherson AJ. Interactions Between the Microbiota and the Immune System. Science (1979) 2012; 1268–1274.

5. Münger E, Montiel-Castro AJ, Langhans W, Pacheco-López G. Reciprocal Interactions Between Gut Microbiota and Host Social Behavior. Front Integr Neurosci 2018; 12: 21.

6. Ezenwa VO, Gerardo NM, Inouye DW, Medina M, Xavier JB. Animal Behavior and the Microbiome. Science (1979) 2012; 338: 198–199.

7. Sherwin E, Bordenstein SR, Quinn JL, Dinan TG, Cryan JF. Microbiota and the social brain. Science (1979) 2019; 366: 587.

8. Diaz Heijtz R, Wang S, Anuar F, Qian Y, Björkholm B, Samuelsson A, et al. Normal gut microbiota modulates brain development and behavior. Proceedings of the National Academy of Sciences 2011; 108: 3047–3052.

9. Neufeld KM, Kang N, Bienenstock J. Reduced anxiety-like behavior and central neurochemical change in germ-free mice. Neurogastroenterology & Motility 2011; 23: 255– 265.

10. Degroote S, Hunting DJ, Baccarelli AA, Takser L. Maternal gut and fetal brain connection: Increased anxiety and reduced social interactions in Wistar rat offspring following peri-conceptional antibiotic exposure. Prog Neuropsychopharmacol Biol Psychiatry 2016; 71: 76–82.

11. Henry LP, Bruijning M, Forsberg SKG, Ayroles JF. The microbiome extends host evolutionary potential. Nat Commun 2021; 12: 5141.

12. Davidson GL, Cooke AC, Johnson CN, Quinn JL. The gut microbiome as a driver of individual variation in cognition and functional behaviour. Philosophical Transactions of the Royal Society B: Biological Sciences 2018; 373: 20170286.

13. Shoji H, Takao K, Hattori S, Miyakawa T. Age-related changes in behavior in C57BL/6J mice from young adulthood to middle age. Mol Brain 2016; 9: 11.

14. Claesson MJ, Jeffery IB, Conde S, Power SE, O’Connor EM, Cusack S, et al. Gut microbiota composition correlates with diet and health in the elderly. Nature 2012; 488: 178–184.

15. Smith P, Willemsen D, Popkes M, Metge F, Gandiwa E, Reichard M, et al. Regulation of life span by the gut microbiota in the short-lived African turquoise killifish. Elife 2017; 6: 1– 26.

16. Liu H, Wang X, Wang HD, Wu J, Ren J, Meng L, et al. Escherichia coli noncoding RNAs can affect gene expression and physiology of Caenorhabditis elegans. Nat Commun 2012; 3: 1–11.

17. Erny D, Prinz M. How microbiota shape microglial phenotypes and epigenetics. Glia 2020; 68: 1655–1672.

18. Canipe LG, Sioda M, Cheatham CL. Diversity of the gut-microbiome related to cognitive behavioral outcomes in healthy older adults. Arch Gerontol Geriatr 2021; 96: 104464.

19. Taniya MA, Chung H-J, Al Mamun A, Alam S, Aziz MdA, Emon NU, et al. Role of Gut Microbiome in Autism Spectrum Disorder and Its Therapeutic Regulation. Front Cell Infect Microbiol 2022; 12.

20. Kelly JR, Minuto C, Cryan JF, Clarke G, Dinan TG. Cross Talk: The Microbiota and Neurodevelopmental Disorders. Front Neurosci 2017; 11.

21. Checa-Ros A, Jeréz-Calero A, Molina-Carballo A, Campoy C, Muñoz-Hoyos A. Current Evidence on the Role of the Gut Microbiome in ADHD Pathophysiology and Therapeutic Implications. Nutrients 2021; 13: 249.

22. Li L, Solvi C, Zhang F, Qi Z, Chittka L, Zhao W. Gut microbiome drives individual memory variation in bumblebees. Nat Commun 2021; 12: 6588.

23. Davidson GL, Wiley N, Cooke AC, Johnson CN, Fouhy F, Reichert MS, et al. Diet induces parallel changes to the gut microbiota and problem solving performance in a wild bird. Sci Rep 2020; 10: 20783.

24. Trevelline BK, Kohl KD. The gut microbiome influences host diet selection behavior. Proceedings of the National Academy of Sciences 2022; 119.

25. Sinotte VM, Renelies-Hamilton J, Taylor BA, Ellegaard KM, Sapountzis P, Vasseur-Cognet M, et al. Synergies Between Division of Labor and Gut Microbiomes of Social Insects. Front Ecol Evol 2020; 7: 1–9.

26. Suenami S, Koto A, Miyazaki R. Basic Structures of Gut Bacterial Communities in Eusocial Insects. Insects 2023; 14: 444.

27. Kwong WK, Moran NA. Gut microbial communities of social bees. Nat Rev Microbiol 2016; 14: 374–384.

28. Robinson GE. Regulation of Division Of Labor In Insect Societies. Annu Rev Entomol 1992; 37: 637–665.

29. Liberti J, Kay T, Quinn A, Kesner L, Frank ET, Cabirol A, et al. The gut microbiota affects the social network of honeybees. Nat Ecol Evol 2022; 6: 1471–1479.

30. Kešnerová L, Emery O, Troilo M, Liberti J, Erkosar B, Engel P. Gut microbiota structure differs between honeybees in winter and summer. ISME J 2020; 14: 801–814.

31. Copeland DC, Maes PW, Mott BM, Anderson KE. Changes in gut microbiota and metabolism associated with phenotypic plasticity in the honey bee Apis mellifera. Front Microbiol 2022; 13.

32. Jones JC, Fruciano C, Marchant J, Hildebrand F, Forslund S, Bork P, et al. The gut microbiome is associated with behavioural task in honey bees. Insectes Soc 2018; 65: 419–429.

33. Gruneck L, Gentekaki E, Khongphinitbunjong K, Popluechai S. Distinct gut microbiota profiles of Asian honey bee (Apis cerana) foragers. Arch Microbiol 2022; 204: 187.

34. Smith CR, Toth AL, Suarez A V., Robinson GE. Genetic and genomic analyses of the division of labour in insect societies. Nat Rev Genet 2008; 9: 735–748.

35. Ben-Shahar Y, Robichon A, Sokolowski MB, Robinson GE. Influence of Gene Action Across Different Time Scales on Behavior. Science (1979) 2002; 296: 741–744.

36. Vernier CL, Krupp JJ, Marcus K, Hefetz A, Levine JD, Ben-Shahar Y. The cuticular hydrocarbon profiles of honey bee workers develop via a socially-modulated innate process. Elife 2019; 8: e41855.

37. Robinson GE, Page RE, Strambi C, Strambi A. Hormonal and Genetic Control of Behavioral Integration in Honey Bee Colonies. Science (1979) 1989; 246: 109–112.

38. Whitfield CW, Cziko A-M, Robinson GE. Gene Expression Profiles in the Brain Predict Behavior in Individual Honey Bees. Science (1979) 2003; 302: 296–299.

39. Ben-Shahar Y, Dudek NL, Robinson GE. Phenotypic deconstruction reveals involvement of manganese transporter malvolio in honey bee division of labor. Journal of Experimental Biology 2004; 207: 3281–3288.

40. Calderone NW, Page RE. Effects of interactions among genotypically diverse nestmates on task specialization by foraging honey bees (Apis mellifera). Behav Ecol Sociobiol 1992; 30: 219–226.

41. Pankiw T, Page Jr RE. Response thresholds to sucrose predict foraging division of labor in honeybees. Behav Ecol Sociobiol 2000; 47: 265–267.

42. Cook CN, Mosqueiro T, Brent CS, Ozturk C, Gadau J, Pinter-Wollman N, et al. Individual differences in learning and biogenic amine levels influence the behavioural division between foraging honeybee scouts and recruits. Journal of Animal Ecology 2019; 88: 236– 246.

43. Tenczar P, Lutz CC, Rao VD, Goldenfeld N, Robinson GE. Automated monitoring reveals extreme interindividual variation and plasticity in honeybee foraging activity levels. Anim Behav 2014; 95: 41–48.

44. Robinson GE. Effects of a juvenile hormone analogue on honey bee foraging behaviour and alarm pheromone production. J Insect Physiol 1985; 31: 277–282.

45. Engel P, James RR, Koga R, Kwong WK, McFrederick QS, Moran NA. Standard methods for research on Apis mellifera gut symbionts. J Apic Res 2013; 52: 1–24.

46. Kešnerová L, Mars RAT, Ellegaard KM, Troilo M, Sauer U, Engel P. Disentangling metabolic functions of bacteria in the honey bee gut. PLoS Biol 2017; 15: e2003467.

47. Leonard SP, Powell JE, Perutka J, Geng P, Heckmann LC, Horak RD, et al. Engineered symbionts activate honey bee immunity and limit pathogens. Science (1979) 2020; 367: 573–576.

48. Powell JE, Martinson VG, Urban-Mead K, Moran NA. Routes of Acquisition of the Gut Microbiota of the Honey Bee Apis mellifera. Appl Environ Microbiol 2014; 80: 7378–7387.

49. Kapheim KM, Rao VD, Yeoman CJ, Wilson BA, White BA, Goldenfeld N, et al. Caste-Specific Differences in Hindgut Microbial Communities of Honey Bees (Apis mellifera). PLoS One 2015; 10: e0123911.

50. Ortiz-Alvarado Y, Clark DR, Vega-Melendez CJ, Flores-Cruz Z, Domingez-Bello MG, Giray T. Antibiotics in hives and their effects on honey bee physiology and behavioral development. Biol Open 2020; 9.

51. Ben-Shahar Y, Thompson CK, Hartz SM, Smith BH, Robinson GE. Differences in performance on a reversal learning test and division of labor in honey bee colonies. Anim Cogn 2000; 3: 119–125.

52. Vernier CL, Chin IM, Adu-Oppong B, Krupp JJ, Levine JD, Dantas G, et al. The gut microbiome defines social group membership in honey bee colonies. Sci Adv 2020; 6: eabd3431.

53. Withers GS, Fahrbach SE, Robinson GE. Effects of experience and juvenile hormone on the organization of the mushroom bodies of honey bees. J Neurobiol 1995; 26: 130–144.

54. Leoncini I, Crauser D, Robinson GE, Le Conte Y. Worker-worker inhibition of honey bee behavioural development independent of queen and brood. Insectes Soc 2004; 51: 392– 394.

55. Caporaso JG, Lauber CL, Walters WA, Berg-Lyons D, Huntley J, Fierer N, et al. Ultra-high-throughput microbial community analysis on the Illumina HiSeq and MiSeq platforms. ISME J 2012; 6: 1621–1624.

56. Callahan BJ, McMurdie PJ, Rosen MJ, Han AW, Johnson AJA, Holmes SP. DADA2: High-resolution sample inference from Illumina amplicon data. Nat Methods 2016; 13: 581–583.

57. Callahan BJ, McMurdie PJ, Holmes SP. Exact sequence variants should replace operational taxonomic units in marker-gene data analysis. ISME J 2017; 11: 2639–2643.

58. Daisley BA, Reid G. BEExact: a Metataxonomic Database Tool for High-Resolution Inference of Bee-Associated Microbial Communities. mSystems 2021; 6.

59. Smith EA, Anderson KE, Corby-Harris V, McFrederick QS, Parish AJ, Rice DW, et al. Reclassification of seven honey bee symbiont strains as Bombella apis. Int J Syst Evol Microbiol 2021; 71.

60. Olofsson TC, Alsterfjord M, Nilson B, Butler È, Vásquez A. Lactobacillus apinorum sp. nov., Lactobacillus mellifer sp. nov., Lactobacillus mellis sp. nov., Lactobacillus melliventris sp. nov., Lactobacillus kimbladii sp. nov., Lactobacillus helsingborgensis sp. nov. and Lactobacillus kullabergensis sp. nov., isolated from the honey stomach of the honeybee Apis mellifera. Int J Syst Evol Microbiol 2014; 64: 3109–3119.

61. Gloor GB, Macklaim JM, Pawlowsky-Glahn V, Egozcue JJ. Microbiome Datasets Are Compositional: And This Is Not Optional. Front Microbiol 2017; 8.

62. Lin H, Peddada S Das. Analysis of compositions of microbiomes with bias correction. Nat Commun 2020; 11: 3514.

63. Aitchison J. The Statistical Analysis of Compositional Data. 1986. Springer Netherlands, Dordrecht.

64. Mandal S, Van Treuren W, White RA, Eggesbø M, Knight R, Peddada SD. Analysis of composition of microbiomes: a novel method for studying microbial composition. Microb Ecol Health Dis 2015; 26.

65. Peddada S, Lin H. Multi-group Analysis of Compositions of Microbiomes with Covariate Adjustments and Repeated Measures. Res Sq 2023.

66. Pfaffl MW. A new mathematical model for relative quantification in real-time RT-PCR. Nucleic Acids Res 2001; 29: 45e–445.

67. Emery O, Schmidt K, Engel P. Immune system stimulation by the gut symbiont Frischella perrara in the honey bee (Apis mellifera). Mol Ecol 2017; 26: 2576–2590.

68. Wu J, Lang H, Mu X, Zhang Z, Su Q, Hu X, et al. Honey bee genetics shape the strain-level structure of gut microbiota in social transmission. Microbiome 2021; 9: 225.

69. Fine JD, Shpigler HY, Ray AM, Beach NJ, Sankey AL, Cash-Ahmed A, et al. Quantifying the effects of pollen nutrition on honey bee queen egg laying with a new laboratory system. PLoS One 2018; 13: 1–16.

70. Galbraith DA, Yang X, Niño EL, Yi S, Grozinger C. Parallel Epigenomic and Transcriptomic Responses to Viral Infection in Honey Bees (Apis mellifera). PLoS Pathog 2015; 11: e1004713.

71. Gernat T, Rao VD, Middendorf M, Dankowicz H, Goldenfeld N, Robinson GE. Automated monitoring of behavior reveals bursty interaction patterns and rapid spreading dynamics in honeybee social networks. Proceedings of the National Academy of Sciences 2018; 115: 1433–1438.

72. Geffre AC, Gernat T, Harwood GP, Jones BM, Morselli Gysi D, Hamilton AR, et al. Honey bee virus causes context-dependent changes in host social behavior. Proceedings of the National Academy of Sciences 2020; 117: 10406–10413.

73. Liaw A, Wiener M. Classification and Regression by randomForest. R News 2002; 2: 18– 22.

74. Hamilton AR, Traniello IM, Ray AM, Caldwell AS, Wickline SA, Robinson GE. Division of labor in honey bees is associated with transcriptional regulatory plasticity in the brain. Journal of Experimental Biology 2019; 222.

75. Klein S, Pasquaretta C, He XJ, Perry C, Søvik E, Devaud J-M, et al. Honey bees increase their foraging performance and frequency of pollen trips through experience. Sci Rep 2019; 9: 6778.

76. R Core Team. R: A language and environment for statistical computing. 2022. R Foundation for Statistical Computing, Vienna, Austria.

77. Pohlert T. The Pairwise Multiple Comparison of Mean Ranks Package (PMCMR). 2014. R package.

78. Oksanen J, Blanchet FG, Friendly M, Kindt R, Legendre P, McGlinn D, et al. vegan: Community Ecology Package. 2017. R package version 2.4-5.

79. Lahti L, Shetty S. microbiome R package. 2019.

80. Martinez Arbizu P. pairwiseAdonis: Pairwise multilevel comparison using adonis. 2020. R package version 0.4.

81. McMurdie PJ, Holmes S. phyloseq: An R Package for Reproducible Interactive Analysis and Graphics of Microbiome Census Data. PLoS One 2013; 8: e61217.

82. Wickham H. Elegant graphics for data analysis. 2016. Springer-Verlag New York.

83. Hervé M. RVAideMemoire: Testing and plotting procedures for biostatistics. 2022. R package.

84. Therneau TM. A package for survival analysis in R. 2022. R package.

85. Kassambara A. rstatix: Pipe-friendly framework for basic statistical test. 2023. R package version 0.7.2.

86. Bates D, Mächler M, Bolker B, Walker S. Fitting Linear Mixed-Effects Models Using lme4. J Stat Softw 2015; 67.

87. Fox J, Weisberg S. An R Companion to Applied Regression, Third Edition. 2018. Sage Publications, Thousand Oaks, CA.

88. Lenth R. emmeans: Estimated marginal means, aka least-squares means. 2023. R package version 1.8.5.

89. Signorell A. DescTools: Tools for descriptive statistics. 2023. R package version 0.99.48.

90. Toth AL, Robinson GE. Worker nutrition and division of labour in honeybees. Anim Behav 2005; 69: 427–435.

91. Toth AL, Kantarovich S, Meisel AF, Robinson GE. Nutritional status influences socially regulated foraging ontogeny in honey bees. Journal of Experimental Biology 2005; 208: 4641–4649.

92. Kubo T, Sasaki M, Nakamura J, Sasagawa H, Ohashi K, Takeuchi H, et al. Change in the Expression of Hypopharyngeal-Gland Proteins of the Worker Honeybees (Apis melliferaL.) with Age and/or Role. J Biochem 1996; 119: 291–295.

93. Ueno T, Takeuchi H, Kawasaki K, Kubo T. Changes in the Gene Expression Profiles of the Hypopharyngeal Gland of Worker Honeybees in Association with Worker Behavior and Hormonal Factors. PLoS One 2015; 10: e0130206.

94. Sigg D, Thompson CM, Mercer AR. Activity-Dependent Changes to the Brain and Behavior of the Honey Bee, *Apis mellifera* (L.). The Journal of Neuroscience 1997; 17: 7148–7156.

95. Fahrbach SE, Robinson GE. Juvenile Hormone, Behavioral Maturation, and Brain Structure in the Honey Bee. Dev Neurosci 1996; 18: 102–114.

96. Withers GS, Fahrbach SE, Robinson GE. Selective neuroanatomical plasticity and division of labour in the honeybee. Nature 1993; 364: 238–240.

97. Finkelstein AB, Brent CS, Giurfa M, Amdam G V. Foraging Experiences Durably Modulate Honey Bees’ Sucrose Responsiveness and Antennal Lobe Biogenic Amine Levels. Sci Rep 2019; 9: 5393.

98. Schulz DJ, Robinson GE. Biogenic amines and division of labor in honey bee colonies: behaviorally related changes in the antennal lobes and age-related changes in the mushroom bodies. J Comp Physiol A 1999; 184: 481–488.

99. Greenberg JK, Xia J, Zhou X, Thatcher SR, Gu X, Ament SA, et al. Behavioral plasticity in honey bees is associated with differences in brain microRNA transcriptome. Genes Brain Behav 2012; 11: 660–670.

100. Sinha S, Jones BM, Traniello IM, Bukhari SA, Halfon MS, Hofmann HA, et al. Behavior-related gene regulatory networks: A new level of organization in the brain. Proceedings of the National Academy of Sciences 2020; 117: 23270–23279.

101. Chandrasekaran S, Ament SA, Eddy JA, Rodriguez-Zas SL, Schatz BR, Price ND, et al. Behavior-specific changes in transcriptional modules lead to distinct and predictable neurogenomic states. Proceedings of the National Academy of Sciences 2011; 108: 18020–18025.

102. Ament SA, Wang Y, Chen C-C, Blatti CA, Hong F, Liang ZS, et al. The Transcription Factor Ultraspiracle Influences Honey Bee Social Behavior and Behavior-Related Gene Expression. PLoS Genet 2012; 8: e1002596.

103. Vellend M. Conceptual Synthesis in Community Ecology. Q Rev Biol 2010; 85: 183–206.

104. Magurran AE. Measuring biological diversity. Current Biology 2021; 31: R1174–R1177.

105. Preston FW. The Commonness, And Rarity, of Species. Ecology 1948; 29: 254–283.

106. Hubbell SP. The Unified Neutral Theory of Biodiversity and Biogeography. 2001. Princeton University Press.

107. Lutz CC, Rodriguez-Zas SL, Fahrbach SE, Robinson GE. Transcriptional response to foraging experience in the honey bee mushroom bodies. Dev Neurobiol 2012; 72: 153– 166.

108. Lutz CC, Robinson GE. Activity-dependent gene expression in honey bee mushroom bodies in response to orientation flight. Journal of Experimental Biology 2013; 216: 2031– 2038.

109. Kapheim KM, Rao VD, Yeoman CJ, Wilson BA, White BA, Goldenfeld N, et al. Caste-Specific Differences in Hindgut Microbial Communities of Honey Bees (Apis mellifera). PLoS One 2015; 10: e0123911.

110. Rueppell O, Bachelier C, Fondrk MK, Page RE. Regulation of life history determines lifespan of worker honey bees (Apis mellifera L.). Exp Gerontol 2007; 42: 1020–1032.

111. Dukas R. Mortality rates of honey bees in the wild. Insectes Soc 2008; 55: 252–255.

112. Visscher PK, Dukas R. Survivorship of foraging honey bees. Insectes Soc 1997; 44: 1–5.

113. Ament SA, Blatti CA, Alaux C, Wheeler MM, Toth AL, Le Conte Y, et al. New meta-analysis tools reveal common transcriptional regulatory basis for multiple determinants of behavior. Proceedings of the National Academy of Sciences 2012; 109: E1801–E1810.

114. Khamis AM, Hamilton AR, Medvedeva YA, Alam T, Alam I, Essack M, et al. Insights into the Transcriptional Architecture of Behavioral Plasticity in the Honey Bee Apis mellifera. Sci Rep 2015; 5: 11136.

115. Anderson KE, Ricigliano VA, Copeland DC, Mott BM, Maes P. Social Interaction is Unnecessary for Hindgut Microbiome Transmission in Honey Bees: The Effect of Diet and Social Exposure on Tissue-Specific Microbiome Assembly. Microb Ecol 2023; 85: 1498– 1513.

116. Ricigliano VA, Anderson KE. Probing the honey bee diet-microbiota-host axis using pollen restriction and organic acid feeding. Insects 2020; 11: 1–14.

