## Supplemental Materials for "Gut microbes contribute to variation in foraging intensity in the honey bee, *Apis mellifera*"

### Supplementary Information

#### Supplementary Figures

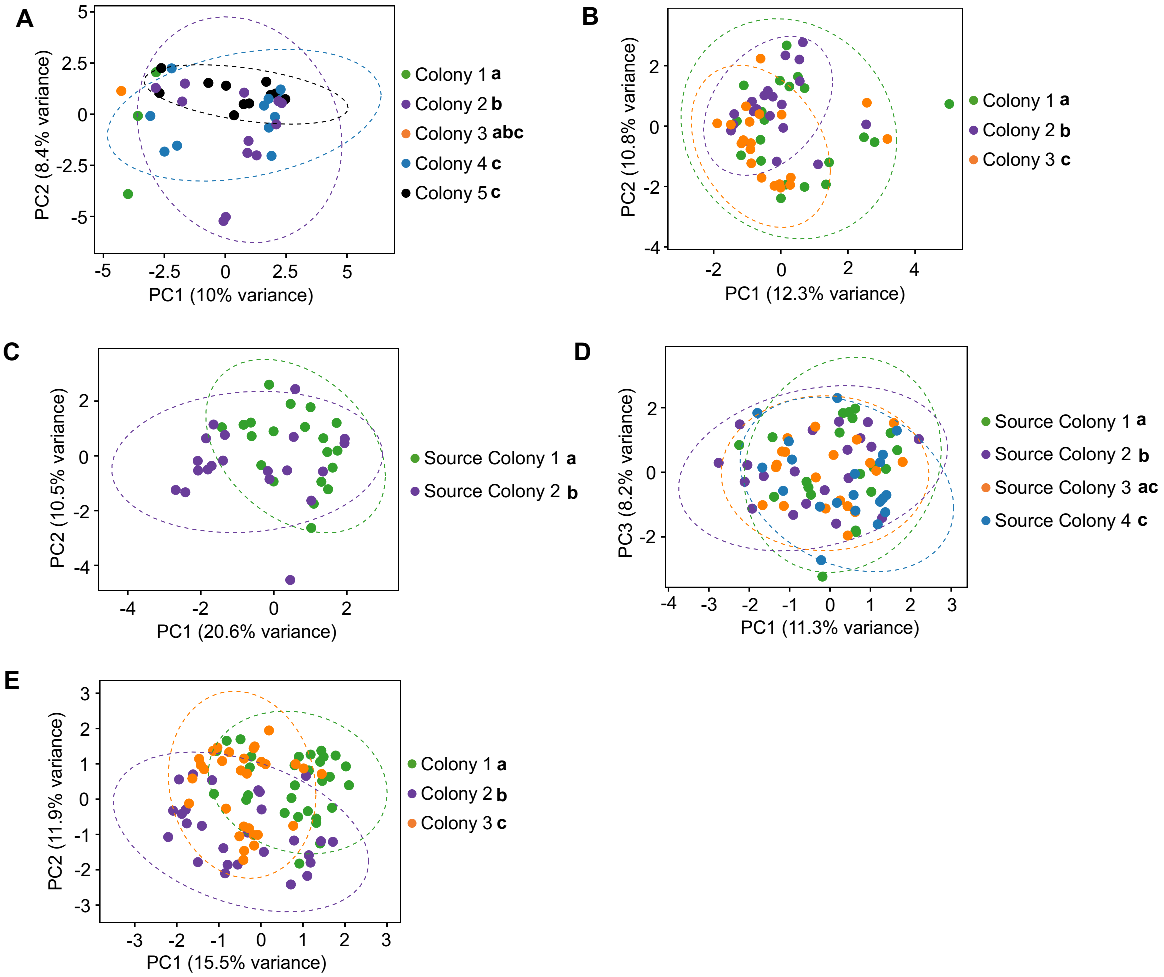


**Supplementary Figure 1. Colonies differ in gut microbial community. (A-B)** Bees from typical colonies differed in gut microbial community. (A) Data reanalyzed from Kapheim et al 2015. Two-way Permutation MANOVA using Aitchison Distance, Task: $F_{1,38} = 1.51$, $R^2 = 0.036$, $p = 0.016$; Colony: $F_{4,38} = 1.72$, $R^2 = 0.165$, $p = 0.001$, Task*Colony: $F_{2,38} = 1.17$, $R^2 = 0.056$, $p = 0.100$. n = 1-12 bees/colony, 5 colonies. (B) New data. Two-way Permutation MANOVA using Aitchison Distance, Task: $F_{1,59} = 4.28$, $R^2 = 0.066$, $p = 0.001$; Colony: $F_{2,59} = 2.31$, $R^2 = 0.071$, $p = 0.001$, Task*Colony: $F_{2,59} = 1.20$, $R^2 = 0.037$, $p = 0.144$. n = 10 bees/colony, 3 colonies. **(C-D)** Bees from different source colonies housed in a single SCC differed in gut microbial community at one week of age (C) and three weeks of age (D). 1 week: Two-way Permutation MANOVA using Aitchison Distance, Task: $F_{1,39} = 0.92$, $R^2 = 0.023$, $p = 0.519$; Source colony: $F_{1,39} = 2.60$, $R^2 = 0.064$, $p = 0.004$, Task*Colony: $F_{1,39} = 1.15$, $R^2 = 0.028$, $p = 0.296$. n = 10 bees/source colony, 2 source colonies. 3 weeks: Two-way Permutation MANOVA using Aitchison, Task: $F_{1,79} = 6.49$, $R^2 = 0.074$, $p = 0.001$; Source colony: $F_{3,79} = 1.82$, $R^2 = 0.062$, $p = 0.001$, Task*Colony: $F_{3,79} = 1.23$, $R^2 = 0.042$, $p = 0.084$. n = 10 bees/source colony, 4 source colonies. **(E)** Bees from different big-back colonies differed in overall gut microbial community. Two-way Permutation MANOVA using Aitchison Distance, Task: $F_{2,86} = 1.84$, $R^2 = 0.038$, $p = 0.011$; Colony: $F_{2,86} = 6.02$, $R^2 = 0.124$, $p = 0.001$, Task*Colony: $F_{4,86} = 0.798$, $R^2 = 0.033$, $p = 0.883$. n = 10 bees/colony, 3 colonies. Depicted as PCA plots. Lowercase letters in legends denote statistically significant groups as determined by Pairwise Permutation MANOVA.

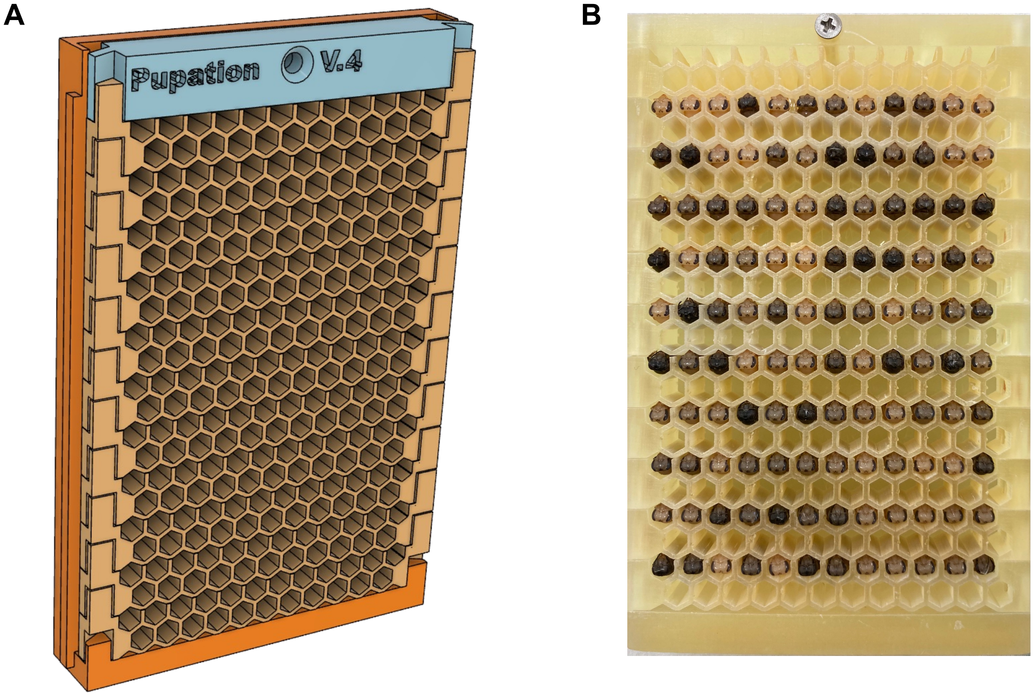


**Supplementary Figure 2. Modular Pupation Plate.** Pupation plates with modular, removeable comb pieces were used for eclosion of bees that lack the dominant honey bee associated gut bacteria. A. Plates were 3D printed with dental grade resin and sterilized through autoclaving and UV irradiation. Full design file found on Mendeley Data (DOI: 10.17632/f2s47y3nhn). B. Photo of pupae in modular pupation plate.

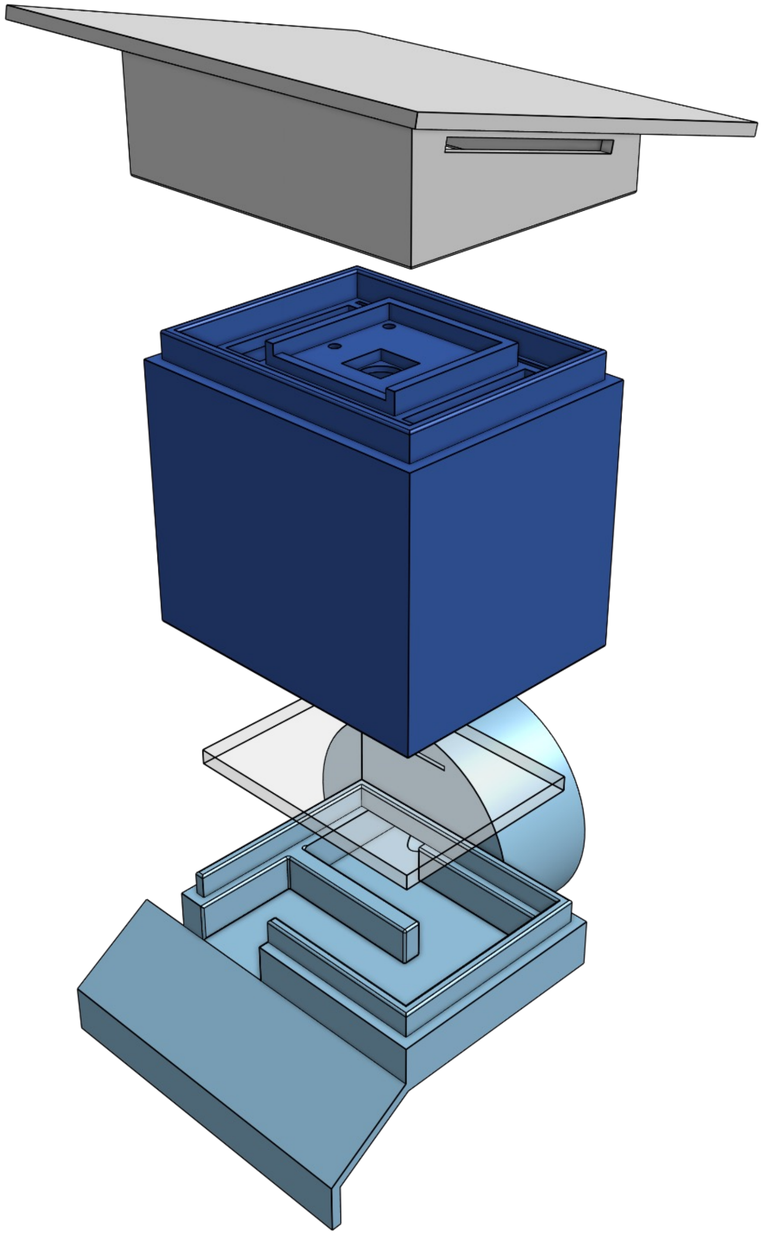

**Supplementary Figure 3. Exploded view of the entrance monitor enclosure.** Shows the base with the landing pad, maze, and connector (light blue), the glass window (semitransparent), the camera mount (dark blue), and the roof that protects the camera from the elements (gray). Camera and camera cable were omitted for visual clarity.

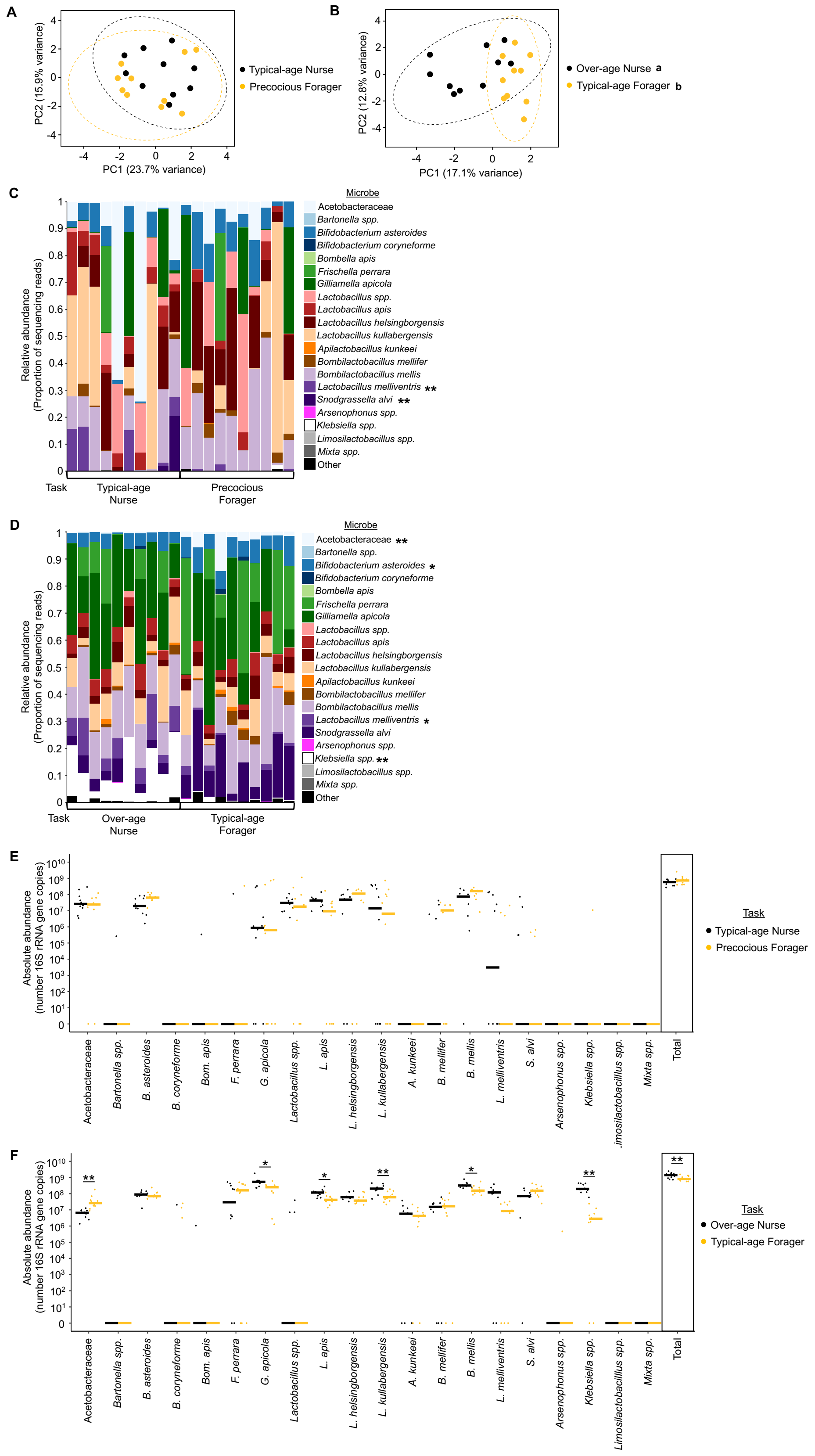

A
PC2 (15.9% variance)
PC1 (23.7% variance)
Typical-age Nurse
Precocious Forager
B
PC2 (12.8% variance)
PC1 (17.1% variance)
Over-age Nurse a
Typical-age Forager b
C
Relative abundance
(Proportion of sequencing reads)
Microbe
Acetobacteraceae
Bartonella spp.
Bifidobacterium asteroides
Bifidobacterium coryneforme
Bombella apis
Frischella perrara
Gilliamella apicola
Lactobacillus spp.
Lactobacillus apis
Lactobacillus helsingborgensis
Lactobacillus kullabergensis
Apilactobacillus kunkeei
Bombilactobacillus mellifer
Bombilactobacillus mellis
Lactobacillus melliventris **
Snodgrassella alvi **
Arsenophonus spp.
Klebsiella spp.
Limosilactobacillus spp.
Mixta spp.
Other
Task
Typical-age Nurse
Precocious Forager
D
Relative abundance
(Proportion of sequencing reads)
Microbe
Acetobacteraceae **
Bartonella spp.
Bifidobacterium asteroides *
Bifidobacterium coryneforme
Bombella apis
Frischella perrara
Gilliamella apicola
Lactobacillus spp.
Lactobacillus apis
Lactobacillus helsingborgensis
Lactobacillus kullabergensis
Apilactobacillus kunkeei
Bombilactobacillus mellifer
Bombilactobacillus mellis
Lactobacillus melliventris *
Snodgrassella alvi
Arsenophonus spp.
Klebsiella spp. **
Limosilactobacillus spp.
Mixta spp.
Other
Task
Over-age Nurse
Typical-age Forager
E
Absolute abundance
(number 16S rRNA gene copies)
Acetobacteraceae
Bartonella spp.
B. asteroides
B. coryneforme
Bom. apis
F. perrara
G. apicola
Lactobacillus spp.
L. apis
L. helsingborgensis
L. kullabergensis
A. kunkeei
B. mellifer
B. mellis
L. melliventris
S. alvi
Arsenophonus spp.
Klebsiella spp.
Limosilactobacillus spp.
Mixta spp.
Total
Task
Typical-age Nurse
Precocious Forager
F
Absolute abundance
(number 16S rRNA gene copies)
Acetobacteraceae
Bartonella spp.
B. asteroides
B. coryneforme
Bom. apis
F. perrara
G. apicola
Lactobacillus spp.
L. apis
L. helsingborgensis
L. kullabergensis
A. kunkeei
B. mellifer
B. mellis
L. melliventris
S. alvi
Arsenophonus spp.
Klebsiella spp.
Limosilactobacillus spp.
Mixta spp.
Total
Task
Over-age Nurse
Typical-age Forager

**Supplementary Figure 4. Bees from single cohort colonies differ in gut microbial community. (A-B)** Age-matched typical-age nurses and precocious foragers from a second single cohort colony did not differ in gut microbial community structure at one week of age (A), but age-matched over-age nurses and typical-age foragers significantly differed in gut microbial community structure at three weeks of age (B). 1 week: Permutation MANOVA using Aitchison Distance, $F_{1,19}$ = 1.36, $R^2$ = 0.07, p = 0.158; n = 10 bees. 3 weeks: Permutation MANOVA using Aitchison Distance, $F_{1,19}$ = 2.64, $R^2$ = 0.128, p = 0.001; n = 10 bees. Depicted as PCA plots. Lowercase letters in legends denote statistically significant groups. **(C-D)** Age-matched typical-age nurses and precocious foragers from a second SCC differed in relative abundance of one individual microbial species (C), and age-matched over-age nurses and typical-age foragers differed in relative abundance of two individual species (D). Depicted as stacked bar plots, with each bar representing a single bee's microbiome. Asterisks in legend: *, $p \leq 0.05$, **, $p \leq 0.01$, ANCOM-BC between nurses and foragers. See Supplementary Table 3 for all p-values. **(E-F)** Age-matched typical-age nurses and precocious foragers from a second SCC did not differ in absolute abundance of individual microbial species or in the total normalized number of 16S rRNA gene copies (E), while age-matched over-age nurses and typical-age foragers differed in absolute abundance of four individual species and the total normalized number of 16S rRNA gene copies (F). $10^x$ number of 16S rRNA gene copies, calculated by multiplying the relative abundance each microbe in each sample (determined through 16S rRNA sequencing) by the normalized number of 16S rRNA gene copies in the sample (determined through qPCR). Depicted as dot plots with all data points plotted, line represents median, n = 10 bees. *, $p \leq 0.05$, **, $p \leq 0.01$, Permutation ANOVA Test between nurses and foragers. See Supplementary Table 3 for all p-values.

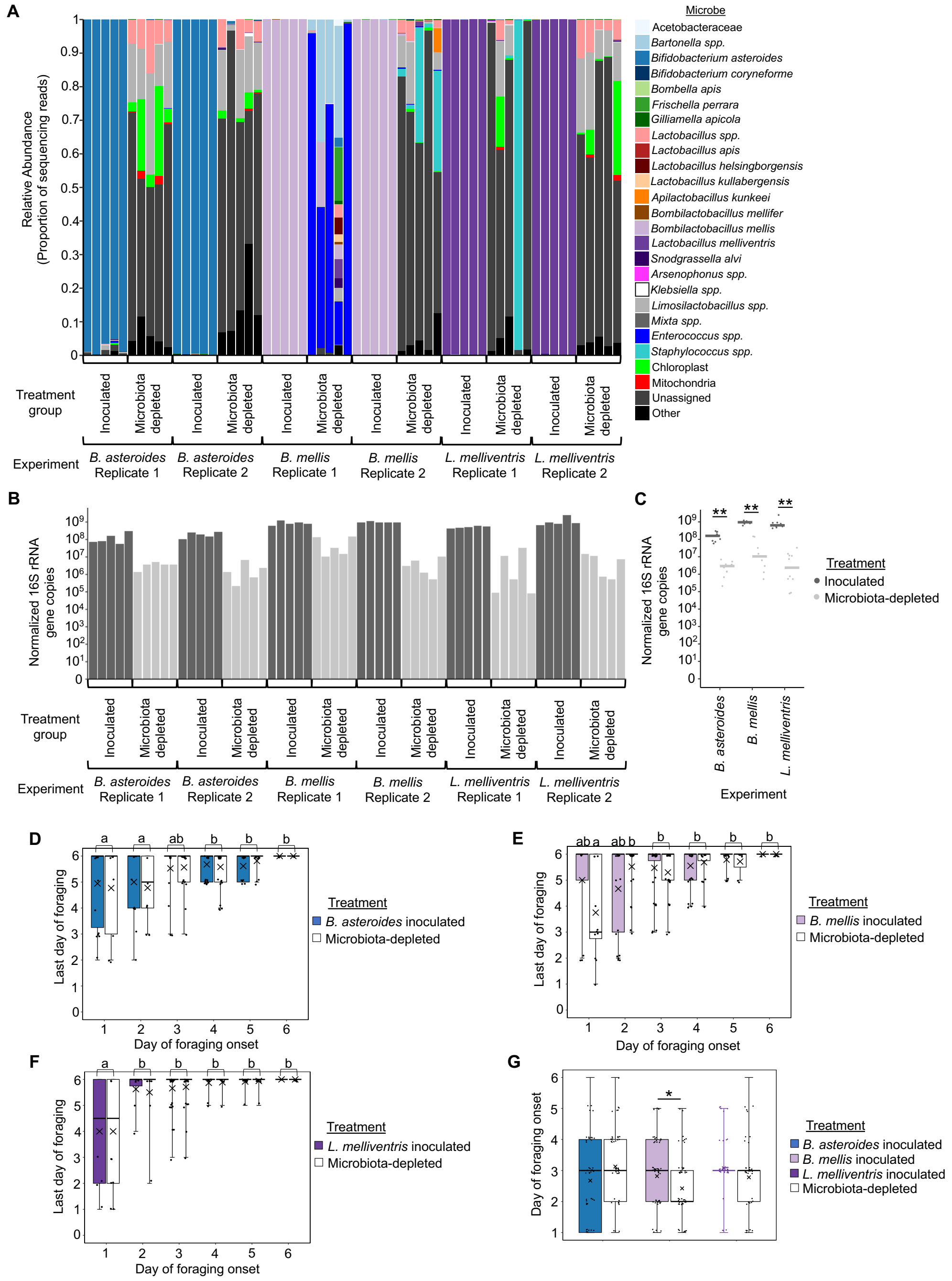


**Supplementary Figure 5. Single microbe inoculated bees differ in gut microbial community. (A)** 16S rRNA sequencing of a subset of single microbe inoculated bee replicates indicated that single microbe inoculated bee guts were mostly composed of the intended honey bee-associated treatment bacteria, while microbiota-depleted bee guts were mostly composed of chloroplast and mitochondrial reads, as well as non-honey bee associated and unassigned bacteria. **(B-C)** Quantitative PCR analysis indicated that single microbe inoculated bee guts had a higher bacterial load than microbiota-depleted bees, measured as absolute abundance of 16S rRNA gene copies normalized to *Actin* gene copies for each sample. *B. asteroides*: Two-way Permutation ANOVA, Treatment: $F_{1,16} = 138.891$, p = 0.001, Replicate: $F_{1,16} = 0.311$, p = 0.590, Treatment*Replicate: $F_{1,16} = 3.917$, p = 0.069. *B. mellis*: Two-way Permutation ANOVA, Treatment: $F_{1,16} = 129.940$, p = 0.001, Replicate: $F_{1,16} = 10.644$, p = 0.010, Treatment*Replicate: $F_{1,16} = 12.944$, p = 0.005. *L. melliventris*: Two-way Permutation ANOVA, Treatment: $F_{1,16} = 66.518$, p = 0.001, Replicate: $F_{1,16} = 1.590$, p = 0.208, Treatment*Replicate: $F_{1,16} = 0.120$, p = 0.725. **(D)** Foragers in *B. asteroides* experimental colonies that began foraging on the first and second days after experimental colony formation had an earlier last day of forage than those who began foraging later, Generalized Linear Mixed Effects Model with log-normal distribution, Treatment: $\chi^2 = 0.081$, p = 0.775, Day of onset: $\chi^2 = 25.35$, p < 0.001, Treatment*Day of onset: $\chi^2 = 0.963$, p = 0.966. **(E)** Microbiota-depleted foragers in *B. mellis* experimental colonies that began foraging on the first day after experimental colony formation had an earlier last day of forage than those who began foraging later, Generalized Linear Mixed Effects Model with log-normal distribution, Treatment: $\chi^2 = 0.036$, p = 0.849, Day of onset: $\chi^2 = 20.68$, p < 0.001, Treatment*Day of onset: $\chi^2 = 14.85$, p = 0.011. **(F)** Foragers in *L. melliventris* experimental colonies that began foraging on the first day after experimental colony formation had an earlier last day of forage than those who began foraging later, Generalized Linear Mixed Effects Model with log-normal distribution, Treatment: $\chi^2 = 0.012$, p = 0.914, Day of onset: $\chi^2 = 51.74$, p < 0.001, Treatment*Day of onset: $\chi^2 = 0.133$, p = 0.999. **(G)** *B. asteroides* and *L. melliventris* inoculated elite foragers began foraging similarly to microbiota-depleted elite foragers, while *B. mellis* inoculated elite foragers began foraging after microbiota-depleted elite foragers. *B. asteroides*: Linear Mixed Effects Model, t-value = 1.657, p = 0.101; *B. mellis*: Generalized Linear Mixed Effects Model, t-value = 2.262, p = 0.024; *L. melliventris*: Linear Mixed Effects Model, t-value = 1.115, p = 0.268. (A), (B) depicted as stacked bar plot. (C) depicted as dot plots with all data points plotted, line represents median. (D), (E), (F), and (G) depicted as box plots with data points plotted, line represents median, x represents mean, and whiskers represent the minimum and maximum values. Lowercase letters in (D-F) denote statistically significantly different groups, with brackets used for those in the same significance group and day of foraging onset for ease of viewing. Asterisks in (G) and (G) denote statistical significance, *, p≤0.05, **, p≤0.001.

### Supplementary Tables and Legends

| Microbe | Nurse relative abundance | Forager relative abundance | Relative: Task adjusted p-value | Nurse absolute abundance | Forager absolute abundance | Absolute: Task adjusted p-value | Absolute: Colony adjusted p-value | Absolute: Task*Colony adjusted p-value |
|---|---|---|---|---|---|---|---|---|
| Acetobacteraceae | 0.050 | 0.063 | 0.787 | 7.42 | 7.68 | **0.030** | 0.482 | 0.361 |
| *Bartonella spp.* | 0.012 | 0.037 | 0.362 | 3.17 | 7.08 | 0.093 | **0.010** | **0.019** |
| *B. asteroides* | 0.080 | 0.064 | 0.350 | 7.86 | 7.81 | 0.727 | 0.573 | 0.444 |
| *B. coryneforme* | 0.014 | 0.011 | 0.322 | 7.10 | 6.89 | 0.666 | 0.727 | 0.109 |
| *Bom. apis* | 0.001 | 0.000 | 0.832 | 0.00 | 0.00 | 0.408 | 0.365 | 0.350 |
| *F. perrara* | 0.045 | 0.049 | 0.889 | 7.56 | 7.66 | 0.843 | **0.014** | 0.421 |
| *G. apicola* | 0.189 | 0.179 | 0.481 | 8.27 | 8.06 | 0.785 | 0.532 | 0.740 |
| *Lactobacillus spp.* | 0.025 | 0.000 | NA | 0.00 | 0.00 | **0.006** | 0.532 | 0.444 |
| *L. apis* | 0.058 | 0.056 | 0.787 | 7.80 | 7.71 | 0.492 | 0.367 | 0.101 |
| *L. helsingborgensis* | 0.074 | 0.018 | **0.001** | 7.73 | 7.05 | 0.065 | 0.482 | 0.869 |
| *L. kullabergensis* | 0.050 | 0.072 | 0.955 | 7.67 | 7.79 | **0.006** | **0.013** | **0.019** |
| *A. kunkeei* | 0.032 | 0.004 | 0.955 | 0.00 | 0.00 | 0.616 | **0.010** | 0.444 |
| *B. mellifer* | 0.011 | 0.024 | 0.481 | 7.01 | 7.36 | 0.232 | 0.732 | 0.444 |
| *B. mellis* | 0.166 | 0.081 | **0.010** | 8.23 | 7.74 | 0.616 | 0.762 | 0.799 |
| *L. melliventris* | 0.071 | 0.016 | **0.000** | 7.81 | 6.93 | **0.019** | 0.532 | 0.846 |
| *S. alvi* | 0.089 | 0.146 | 0.787 | 7.94 | 8.06 | **0.041** | 0.762 | 0.444 |
| *Arsenophonus spp.* | 0.001 | 0.138 | 0.115 | 0.00 | 0.00 | 0.408 | 0.668 | 0.460 |
| *Klebsiella spp.* | 0.019 | 0.000 | NA | 3.11 | 0.00 | **0.006** | 0.532 | 0.444 |
| *Limosilactobacillus spp.* | 0.000 | 0.000 | NA | 0.00 | 0.00 | NA | NA | NA |
| *Mixta spp.* | 0.001 | 0.001 | NA | 0.00 | 0.00 | 0.843 | 0.727 | 0.804 |
| Total | NA | NA | NA | 9.005 | 8.935 | 0.202 | 0.671 | 0.618 |

**Supplementary Table 1: Abundance of each microbe in the gut microbial communities of nurse and forager bees.** Relative abundance, mean proportion of 16S rRNA sequencing reads per sample. Absolute abundance, $10^x$ median number of 16S rRNA gene copies per sample. This was calculated by multiplying the relative abundance of each microbe in each sample (determined through 16S rRNA sequencing) by the normalized number of 16S rRNA gene copies in the sample (determined through qPCR). Relative abundances were analyzed via ANCOM-BC, while absolute abundances were analyzed via Permutation ANOVA. p-values were adjusted to account for multiple comparisons through FDR adjustment. NAs represent taxa that were not present in enough samples to reliably analyze. n = 10 bees/task/colony, 3 colonies.

| Age | Microbe | Nurse relative abundance | Forager relative abundance | Relative: Task adjusted p-value | Nurse absolute abundance | Forager absolute abundance | Absolute: Task adjusted p-value | Absolute: Source colony adjusted p-value | Absolute: Task*Source adjusted p-value |
|---|---|---|---|---|---|---|---|---|---|
| 1 week | Acetobacteraceae | 0.068 | 0.162 | 0.852 | 7.46 | 7.58 | 0.390 | **0.004** | 0.667 |
| 1 week | *Bartonella spp.* | 0.004 | 0.003 | 0.852 | 0.00 | 0.00 | 0.711 | 0.608 | 0.953 |
| 1 week | *B. asteroides* | 0.059 | 0.060 | 0.947 | 7.73 | 7.51 | 0.390 | **0.006** | 0.602 |
| 1 week | *B. coryneforme* | 0.002 | 0.002 | 0.889 | 0.00 | 0.00 | 0.882 | 0.062 | 0.997 |
| 1 week | *Bom. apis* | 0.000 | 0.000 | NA | 0.00 | 0.00 | NA | NA | NA |
| 1 week | *F. perrara* | 0.093 | 0.061 | 0.852 | 6.43 | 0.00 | 0.711 | 0.608 | 0.667 |
| 1 week | *G. apicola* | 0.293 | 0.168 | 0.852 | 8.35 | 7.99 | 0.390 | **0.004** | 0.683 |
| 1 week | *Lactobacillus spp.* | 0.066 | 0.057 | 0.852 | 7.53 | 6.46 | 0.390 | 0.219 | 0.400 |
| 1 week | *L. apis* | 0.044 | 0.040 | 0.852 | 7.65 | 7.40 | 0.711 | **0.030** | 0.859 |
| 1 week | *L. helsingborgensis* | 0.063 | 0.071 | 0.852 | 7.49 | 7.49 | 0.698 | 0.401 | 0.799 |
| 1 week | *L. kullabergensis* | 0.054 | 0.054 | 0.852 | 0.00 | 7.24 | 0.649 | 0.656 | 0.667 |
| 1 week | *A. kunkeei* | 0.000 | 0.000 | NA | 0.00 | 0.00 | NA | NA | NA |
| 1 week | *B. mellifer* | 0.018 | 0.019 | 0.852 | 7.18 | 6.85 | 0.649 | **0.004** | 0.953 |
| 1 week | *B. mellis* | 0.133 | 0.133 | 0.852 | 8.17 | 7.84 | 0.390 | **0.004** | 0.667 |
| 1 week | *L. melliventris* | 0.042 | 0.020 | 0.852 | 3.49 | 0.00 | 0.649 | 0.107 | 0.405 |
| 1 week | *S. alvi* | 0.063 | 0.092 | 0.852 | 7.71 | 7.61 | 0.600 | **0.016** | 0.444 |
| 1 week | *Arsenophonus spp.* | 0.000 | 0.000 | NA | 0.00 | 0.00 | NA | NA | NA |
| 1 week | *Klebsiella spp.* | 0.000 | 0.000 | NA | 0.00 | 0.00 | 0.711 | 0.480 | 0.739 |
| 1 week | *Limosilactobacillus spp.* | 0.000 | 0.000 | NA | 0.00 | 0.00 | NA | NA | NA |
| 1 week | *Mixta spp.* | 0.000 | 0.057 | NA | 0.00 | 0.00 | 0.711 | 0.656 | 0.405 |
| 1 week | Total | NA | NA | NA | 8.892 | 8.78 | **0.001** | **0.001** | 0.650 |
| 3 weeks | Acetobacteraceae | 0.004 | 0.012 | **0.000** | 6.54 | 6.95 | 0.065 | 0.543 | 0.295 |
| 3 weeks | *Bartonella spp.* | 0.002 | 0.028 | **0.009** | 0.00 | 0.00 | 0.111 | **0.019** | 0.604 |
| 3 weeks | *B. asteroides* | 0.056 | 0.122 | **0.050** | 8.01 | 7.98 | 0.847 | 0.997 | 0.740 |
| 3 weeks | *B. coryneforme* | 0.009 | 0.007 | 0.296 | 7.12 | 0.00 | **0.003** | 0.068 | 0.740 |
| 3 weeks | *Bom. apis* | 0.000 | 0.000 | NA | 0.00 | 0.00 | 0.320 | 0.899 | 0.740 |
| 3 weeks | *F. perrara* | 0.062 | 0.145 | **0.012** | 7.88 | 7.04 | 0.065 | 0.630 | 0.740 |
| 3 weeks | *G. apicola* | 0.153 | 0.175 | 0.778 | 8.35 | 8.14 | **0.038** | 0.363 | 0.698 |
| 3 weeks | *Lactobacillus spp.* | 0.002 | 0.001 | 0.647 | 0.00 | 0.00 | 0.354 | 0.345 | 0.701 |
| 3 weeks | *L. apis* | 0.057 | 0.054 | 0.916 | 7.97 | 7.59 | 0.949 | 0.136 | 0.701 |
| 3 weeks | *L. helsingborgensis* | 0.051 | 0.057 | 0.777 | 7.89 | 7.64 | 0.717 | 0.931 | 0.740 |
| 3 weeks | *L. kullabergensis* | 0.046 | 0.046 | 0.706 | 7.84 | 7.51 | 0.065 | 0.068 | 0.740 |
| 3 weeks | *A. kunkeei* | 0.000 | 0.000 | NA | 0.00 | 0.00 | 0.607 | 0.630 | 0.740 |
| 3 weeks | *B. mellifer* | 0.010 | 0.020 | 0.069 | 7.26 | 7.23 | 0.624 | 0.630 | 0.907 |
| 3 weeks | *B. mellis* | 0.107 | 0.072 | 0.069 | 8.25 | 7.68 | **0.003** | **0.019** | 0.527 |
| 3 weeks | *L. melliventris* | 0.044 | 0.013 | **0.003** | 7.84 | 6.93 | **0.003** | 0.551 | 0.637 |
| 3 weeks | *S. alvi* | 0.075 | 0.215 | **0.000** | 7.97 | 8.24 | **0.003** | 0.068 | 0.295 |
| 3 weeks | *Arsenophonus spp.* | 0.000 | 0.020 | NA | 0.00 | 0.00 | 0.757 | 0.997 | 0.527 |
| 3 weeks | *Klebsiella spp.* | 0.314 | 0.011 | **0.000** | 8.64 | 6.53 | **0.003** | 0.899 | 0.875 |
| 3 weeks | *Limosilactobacillus spp.* | 0.000 | 0.000 | NA | 0.00 | 0.00 | NA | NA | NA |
| 3 weeks | *Mixta spp.* | 0.002 | 0.000 | NA | 0.00 | 0.00 | **0.003** | 0.630 | 0.740 |
| 3 weeks | Total | NA | NA | NA | 9.226 | 8.925 | **0.001** | **0.017** | 0.123 |

**Supplementary Table 2: Abundance of each microbe in the gut microbial communities of nurse and forager bees from a single cohort colony.** Relative abundance, mean proportion of 16S rRNA sequencing reads per sample. Absolute abundance, $10^x$ median number of 16S rRNA gene copies per sample. This was calculated by multiplying the relative abundance of each microbe in each sample (determined through 16S rRNA sequencing) by the normalized number of 16S rRNA gene copies in the sample (determined through qPCR). Relative abundances were analyzed via ANCOM-BC, while absolute abundances were analyzed via Permutation ANOVA. p-values were adjusted to account for multiple comparisons through FDR adjustment. NAs represent taxa that were not present in enough samples to reliably analyze. n = 10 bees/task/source colony at each age, 3 source colonies.

| Age | Microbe | Nurse relative abundance | Forager relative abundance | Relative: Task adjusted p-value | Nurse absolute abundance | Forager absolute abundance | Absolute: Task adjusted p-value |
|---|---|---|---|---|---|---|---|
| 1 week | Acetobacteraceae | 0.187 | 0.052 | 0.591 | 7.41 | 7.38 | 0.457 |
| 1 week | *Bartonella spp.* | 0.000 | 0.000 | NA | 0.00 | 0.00 | 0.710 |
| 1 week | *B. asteroides* | 0.053 | 0.101 | 0.869 | 7.28 | 7.80 | 0.200 |
| 1 week | *B. coryneforme* | 0.000 | 0.000 | NA | 0.00 | 0.00 | NA |
| 1 week | *Bom. apis* | 0.000 | 0.000 | NA | 0.00 | 0.00 | 0.200 |
| 1 week | *F. perrara* | 0.032 | 0.040 | NA | 0.00 | 0.00 | 0.977 |
| 1 week | *G. apicola* | 0.073 | 0.129 | 0.869 | 5.94 | 5.80 | 0.953 |
| 1 week | *Lactobacillus spp.* | 0.081 | 0.110 | 0.869 | 7.48 | 7.25 | 0.386 |
| 1 week | *L. apis* | 0.073 | 0.024 | 0.126 | 7.62 | 6.96 | 0.200 |
| 1 week | *L. helsingborgensis* | 0.094 | 0.176 | 0.869 | 7.68 | 8.05 | 0.557 |
| 1 week | *L. kullabergensis* | 0.202 | 0.139 | 0.591 | 7.14 | 6.82 | 0.953 |
| 1 week | *A. kunkeei* | 0.000 | 0.000 | NA | 0.00 | 0.00 | NA |
| 1 week | *B. mellifer* | 0.009 | 0.020 | 0.869 | 0.00 | 7.02 | 0.386 |
| 1 week | *B. mellis* | 0.117 | 0.203 | 0.869 | 7.87 | 8.21 | 0.304 |
| 1 week | *L. melliventris* | 0.055 | 0.004 | **0.022** | 3.49 | 0.00 | 0.386 |
| 1 week | *S. alvi* | 0.023 | 0.000 | **0.022** | 0.00 | 0.00 | 0.650 |
| 1 week | *Arsenophonus spp.* | 0.000 | 0.000 | NA | 0.00 | 0.00 | NA |
| 1 week | *Klebsiella spp.* | 0.000 | 0.001 | NA | 0.00 | 0.00 | 0.386 |
| 1 week | *Limosilactobacillus spp.* | 0.000 | 0.000 | NA | 0.00 | 0.00 | NA |
| 1 week | *Mixta spp.* | 0.000 | 0.000 | NA | 0.00 | 0.00 | NA |
| 1 week | Total | NA | NA | NA | 8.767 | 8.865 | 0.218 |
| 3 weeks | Acetobacteraceae | 0.003 | 0.036 | **0.001** | 6.82 | 7.42 | **0.006** |
| 3 weeks | *Bartonella spp.* | 0.000 | 0.000 | NA | 0.00 | 0.00 | NA |
| 3 weeks | *B. asteroides* | 0.044 | 0.072 | **0.042** | 7.95 | 7.85 | 1.000 |
| 3 weeks | *B. coryneforme* | 0.001 | 0.004 | NA | 0.00 | 0.00 | 0.569 |
| 3 weeks | *Bom. apis* | 0.000 | 0.000 | NA | 0.00 | 0.00 | 0.908 |
| 3 weeks | *F. perrara* | 0.069 | 0.180 | 0.205 | 7.47 | 8.21 | 1.000 |
| 3 weeks | *G. apicola* | 0.254 | 0.197 | 0.427 | 8.73 | 8.40 | **0.027** |
| 3 weeks | *Lactobacillus spp.* | 0.003 | 0.000 | NA | 0.00 | 0.00 | 0.344 |
| 3 weeks | *L. apis* | 0.056 | 0.040 | 0.739 | 8.07 | 7.62 | **0.018** |
| 3 weeks | *L. helsingborgensis* | 0.038 | 0.042 | 0.667 | 7.78 | 7.57 | 0.306 |
| 3 weeks | *L. kullabergensis* | 0.106 | 0.067 | 0.675 | 8.31 | 7.78 | **0.006** |
| 3 weeks | *A. kunkeei* | 0.004 | 0.005 | 0.986 | 6.76 | 6.63 | 0.779 |
| 3 weeks | *B. mellifer* | 0.009 | 0.021 | 0.362 | 7.19 | 7.22 | 0.779 |
| 3 weeks | *B. mellis* | 0.172 | 0.162 | 0.985 | 8.50 | 8.19 | **0.025** |
| 3 weeks | *L. melliventris* | 0.068 | 0.010 | **0.033** | 8.07 | 6.94 | 0.497 |
| 3 weeks | *S. alvi* | 0.038 | 0.148 | 0.051 | 7.85 | 8.18 | 0.159 |
| 3 weeks | *Arsenophonus spp.* | 0.000 | 0.000 | NA | 0.00 | 0.00 | 0.344 |
| 3 weeks | *Klebsiella spp.* | 0.128 | 0.005 | **0.001** | 8.30 | 6.45 | **0.006** |
| 3 weeks | *Limosilactobacillus spp.* | 0.000 | 0.000 | NA | 0.00 | 0.00 | NA |
| 3 weeks | *Mixta spp.* | 0.000 | 0.000 | NA | 0.00 | 0.00 | NA |
| 3 weeks | Total | NA | NA | NA | 9.273 | 9.072 | **0.001** |

**Supplementary Table 3: Abundance of each microbe in the gut microbial communities of nurse and forager bees from a single cohort colony.** Relative abundance, mean proportion of 16S rRNA sequencing reads per sample. Absolute abundance, $10^x$ median number of 16S rRNA gene copies per sample. This was calculated by multiplying the relative abundance of each microbe in each sample (determined through 16S rRNA sequencing) by the normalized number of 16S rRNA gene copies in the sample (determined through qPCR). Relative abundances were analyzed via ANCOM-BC, while absolute abundances were analyzed via Permutation ANOVA. p-values were adjusted to account for multiple comparisons through FDR adjustment. NAs represent taxa that were not present in enough samples to reliably analyze. n = 10 bees/task at each age, 1 source colony.

| Microbe | Inactive forager relative abundance | Active forager relative abundance | Relative: Task adjusted p-value | Inactive forager absolute abundance | Active forager absolute abundance | Absolute: Task adjusted p-value | Absolute: Source colony adjusted p-value | Absolute: Task*Source adjusted p-value |
|---|---|---|---|---|---|---|---|---|
| Acetobacteraceae | 0.121 | 0.037 | 0.180 | 7.67 | 7.04 | 0.573 | **0.003** | 0.351 |
| *Bartonella spp.* | 0.002 | 0.000 | NA | 0.00 | 0.00 | 0.371 | **0.045** | 0.351 |
| *B. asteroides* | 0.078 | 0.124 | 0.757 | 7.81 | 7.70 | 0.618 | **0.046** | 0.568 |
| *B. coryneforme* | 0.002 | 0.002 | NA | 0.00 | 0.00 | 0.835 | 0.860 | 0.645 |
| *Bom. apis* | 0.000 | 0.000 | NA | 0.00 | 0.00 | 0.835 | 0.729 | 0.389 |
| *F. perrara* | 0.082 | 0.183 | 0.481 | 6.36 | 7.60 | 0.860 | 0.483 | 0.645 |
| *G. apicola* | 0.185 | 0.161 | 0.646 | 7.90 | 7.71 | 0.371 | **0.003** | 0.645 |
| *Lactobacillus spp.* | 0.144 | 0.016 | **0.004** | 7.74 | 6.08 | **0.017** | 0.647 | 0.568 |
| *L. apis* | 0.087 | 0.018 | **0.003** | 7.41 | 6.73 | 0.860 | **0.003** | 0.546 |
| *L. helsingborgensis* | 0.116 | 0.070 | 0.125 | 7.86 | 7.50 | 0.835 | 0.158 | 0.990 |
| *L. kullabergensis* | 0.018 | 0.043 | 0.481 | 0.00 | 3.58 | 0.391 | **0.027** | 0.351 |
| *A. kunkeei* | 0.000 | 0.001 | NA | 0.00 | 0.00 | 0.264 | **0.009** | 0.351 |
| *B. mellifer* | 0.004 | 0.017 | 0.757 | 0.00 | 3.26 | 0.264 | **0.003** | 0.351 |
| *B. mellis* | 0.047 | 0.255 | **0.014** | 7.36 | 8.05 | 0.391 | **0.043** | 0.990 |
| *L. melliventris* | 0.070 | 0.035 | 0.180 | 7.47 | 6.85 | 0.423 | **0.003** | 0.990 |
| *S. alvi* | 0.038 | 0.037 | 0.536 | 7.19 | 7.00 | 0.541 | 0.323 | 0.990 |
| *Arsenophonus spp.* | 0.000 | 0.000 | NA | 0.00 | 0.00 | NA | NA | NA |
| *Klebsiella spp.* | 0.005 | 0.000 | NA | 0.00 | 0.00 | 0.136 | 0.163 | 0.351 |
| *Limosilactobacillus spp* | 0.000 | 0.000 | NA | 0.00 | 0.00 | NA | NA | NA |
| *Mixta spp.* | 0.000 | 0.000 | NA | 0.00 | 0.00 | NA | NA | NA |
| Total | NA | NA | NA | 8.959 | 8.658 | **0.001** | 0.015 | 0.465 |

**Supplementary Table 4: Abundance of each microbe in the gut microbial communities of inactive and active forager bees from big-back colonies.** Relative abundance, mean proportion of 16S rRNA sequencing reads per sample. Absolute abundance, $10^x$ median number of 16S rRNA gene copies per sample. This was calculated by multiplying the relative abundance of each microbe in each sample (determined through 16S rRNA sequencing) by the normalized number of 16S rRNA gene copies in the sample (determined through qPCR). Relative abundances were analyzed via ANCOM-BC, while absolute abundances were analyzed via Permutation ANOVA. p-values were adjusted to account for multiple comparisons through FDR adjustment. NAs represent taxa that were not present in enough samples to reliably analyze. n = 10 bees/task/colony, 3 colonies.

| Fig. | Test microbe | Measure | Day | Statistical test | | Comparison | Test statistic | Test statistic value | p-value |
|---|---|---|---|---|---|---|---|---|---|
| 4A | *B. asteroides* | Cumulative proportion bees foraging per treatment group | All | Cox Proportional Hazards | | Treatment | z | -0.331 | 0.741 |
| | | | | **mean inoculated group** | **mean depleted group** | | | | |
| | | | 1 | 0.068 | 0.041 | | | | |
| | | | 2 | 0.123 | 0.075 | | | | |
| | | | 3 | 0.188 | 0.148 | | | | |
| | | | 4 | 0.288 | 0.253 | | | | |
| | | | 5 | 0.329 | 0.304 | | | | |
| | | | 6 | 0.343 | 0.322 | | | | |
| 4B | *B. asteroides* | Proportion foraging events per treatment group | All | Linear Mixed Effects Model | | Treatment | F(1,33) | 21.165 | **<0.001** |
| | | | | | | Day | F(5,33) | 0 | 1 |
| | | | | | | Treatment*Day | F(5,33) | 1.987 | **0.107** |
| | | | | Pairwise (emmeans posthoc) | | | | | |
| | | | | **mean inoculated group** | **mean depleted group** | | | | |
| | | | 1 | 0.708 | 0.292 | Treatment | t-ratio | 4.004 | **<0.001** |
| | | | 2 | 0.664 | 0.336 | Treatment | t-ratio | 3.146 | **0.004** |
| | | | 3 | 0.586 | 0.414 | Treatment | t-ratio | 1.656 | 0.107 |
| | | | 4 | 0.517 | 0.483 | Treatment | t-ratio | 0.318 | 0.753 |
| | | | 5 | 0.55 | 0.45 | Treatment | t-ratio | 0.963 | 0.343 |
| | | | 6 | 0.561 | 0.439 | Treatment | t-ratio | 1.182 | 0.246 |
| 4C | *B. asteroides* | Number of foragers per treatment group | All | Linear Mixed Effects Model | | Treatment | F(1,33) | 0.409 | 0.527 |
| | | | | | | Day | F(5,33) | 12.095 | **<0.001** |
| | | | | | | Treatment*Day | F(5,33) | 0.14 | 0.982 |
| | | | | Pairwise (emmeans posthoc) | | | | | |
| | | | | **mean inoculated group** | **mean depleted group** | | | | |
| | | | 1 | 4.5 | 3 | Treatment | t-ratio | 0.409 | 0.685 |
| | | | 2 | 8 | 5 | Treatment | t-ratio | 0.817 | 0.42 |
| | | | 3 | 12.2 | 10.5 | Treatment | t-ratio | 0.477 | 0.637 |
| | | | 4 | 17.5 | 17.2 | Treatment | t-ratio | 0.068 | 0.946 |
| | | | 5 | 19.5 | 19.5 | Treatment | t-ratio | 0 | 1 |
| | | | 6 | 16.2 | 17 | Treatment | t-ratio | -0.204 | 0.839 |
| 4D | *B. asteroides* | Proportion foraging events per individual | All | Generalized Linear Mixed Effects Model | | Treatment | $\chi^2$ | 4.985 | **0.026** |
| | | | | | | Day | $\chi^2$ | 382.933 | **<0.001** |
| | | | | | | Treatment*Day | $\chi^2$ | 26.315 | **<0.001** |
| | | | | Pairwise (emmeans posthoc) | | | | | |
| | | | | **mean inoculated group** | **mean depleted group** | | | | |
| | | | 1 | 0.157 | 0.09 | Treatment | z-ratio | 5.403 | **<0.001** |
| | | | 2 | 0.083 | 0.067 | Treatment | z-ratio | 1.691 | 0.091 |

| | | | | | | | | | |
|---|---|---|---|---|---|---|---|---|---|
| | | | 3 | 0.048 | 0.039 | Treatment | z-ratio | 0.87 | 0.384 |
| | | | 4 | 0.03 | 0.028 | Treatment | z-ratio | -0.026 | 0.979 |
| | | | 5 | 0.028 | 0.023 | Treatment | z-ratio | 0.691 | 0.49 |
| | | | 6 | 0.035 | 0.026 | Treatment | z-ratio | 1.318 | 0.187 |
| 4E | *B. asteroides* | Proportion of elite foragers per treatment group | All | Linear Mixed Effects Model | | Treatment | F(1,33) | 15.799 | **<0.001** |
| | | | | | | Day | F(5,33) | 0 | 1 |
| | | | | | | Treatment*Day | F(5,33) | 3.151 | **0.02** |
| | | | | Pairwise (emmeans posthoc) | | | | | |
| | | | | **mean inoculated group** | **mean depleted group** | | | | |
| | | | 1 | 0.792 | 0.208 | Treatment | t-ratio | 4.039 | **<0.001** |
| | | | 2 | 0.762 | 0.238 | Treatment | t-ratio | 3.635 | **<0.001** |
| | | | 3 | 0.586 | 0.414 | Treatment | t-ratio | 1.187 | 0.244 |
| | | | 4 | 0.492 | 0.508 | Treatment | t-ratio | -0.113 | 0.911 |
| | | | 5 | 0.554 | 0.446 | Treatment | t-ratio | 0.742 | 0.464 |
| | | | 6 | 0.518 | 0.482 | Treatment | t-ratio | 0.247 | 0.806 |
| 5A | *B. mellis* | Cumulative proportion bees foraging per treatment group | All | Cox Proportional Hazards | | Treatment | z | 1.525 | 0.127 |
| | | | | **mean inoculated group** | **mean depleted group** | | | | |
| | | | 1 | 0.048 | 0.051 | | | | |
| | | | 2 | 0.134 | 0.155 | | | | |
| | | | 3 | 0.269 | 0.256 | | | | |
| | | | 4 | 0.374 | 0.316 | | | | |
| | | | 5 | 0.425 | 0.34 | | | | |
| | | | 6 | 0.435 | 0.351 | | | | |
| 5B | *B. mellis* | Proportion foraging events per treatment group | All | Linear Mixed Effects Model | | Treatment | F(1,29.091) | 0.036 | 0.851 |
| | | | | | | Day | F(5,29.719) | 0 | 1 |
| | | | | | | Treatment*Day | F(5,29.091) | 1.981 | 0.111 |
| | | | | Pairwise (emmeans posthoc) | | | | | |
| | | | | **mean inoculated group** | **mean depleted group** | | | | |
| | | | 1 | 0.418 | 0.582 | Treatment | t-ratio | 1.307 | 0.201 |
| | | | 2 | 0.39 | 0.61 | Treatment | t-ratio | 1.757 | 0.089 |
| | | | 3 | 0.439 | 0.561 | Treatment | t-ratio | 1.126 | 0.269 |
| | | | 4 | 0.554 | 0.446 | Treatment | t-ratio | -0.995 | 0.328 |
| | | | 5 | 0.549 | 0.451 | Treatment | t-ratio | -0.898 | 0.376 |
| | | | 6 | 0.578 | 0.422 | Treatment | t-ratio | -1.442 | 0.16 |
| 5C | *B. mellis* | Number of foragers per treatment group | All | Linear Mixed Effects Model | | Treatment | F(1,33) | 3.183 | 0.084 |
| | | | | | | Day | F(5,33) | 14.918 | **<0.001** |
| | | | | | | Treatment*Day | F(5,33) | 0.765 | 0.582 |
| | | | | Pairwise (emmeans posthoc) | | | | | |
| | | | | **mean inoculated group** | **mean depleted group** | | | | |
| | | | 1 | 3 | 3 | Treatment | t-ratio | 0 | 1 |

| | | | | | | | | | |
|---|---|---|---|---|---|---|---|---|---|
| | | | 2 | 8.25 | 8.75 | Treatment | t-ratio | 0.143 | 0.887 |
| | | | 3 | 15.2 | 15.2 | Treatment | t-ratio | 0 | 1 |
| | | | 4 | 21.2 | 17.2 | Treatment | t-ratio | -1.146 | 0.26 |
| | | | 5 | 22.8 | 17 | Treatment | t-ratio | -1.648 | 0.109 |
| | | | 6 | 20.5 | 14.5 | Treatment | t-ratio | -1.719 | 0.095 |
| 5D | *B. mellis* | Proportion foraging events per individual | All | Generalized Linear Mixed Effects Model | | Treatment | $\chi^2$ | 3.626 | 0.057 |
| | | | | | | Day | $\chi^2$ | 255.497 | **<0.001** |
| | | | | | | Treatment*Day | $\chi^2$ | 6.552 | 0.256 |
| | | | | Pairwise (emmeans posthoc) | | | | | |
| | | | | **mean inoculated group** | **mean depleted group** | | | | |
| | | | 1 | 0.105 | 0.145 | Treatment | z-ratio | 2.313 | **0.021** |
| | | | 2 | 0.036 | 0.052 | Treatment | z-ratio | 2.194 | **0.028** |
| | | | 3 | 0.029 | 0.037 | Treatment | z-ratio | 1.487 | 0.137 |
| | | | 4 | 0.026 | 0.026 | Treatment | z-ratio | 0.177 | 0.859 |
| | | | 5 | 0.024 | 0.027 | Treatment | z-ratio | 1.026 | 0.305 |
| | | | 6 | 0.028 | 0.029 | Treatment | z-ratio | 0.792 | 0.429 |
| 5E | *B. mellis* | Proportion of elite foragers per treatment group | All | Linear Mixed Effects Model | | Treatment | F(1,29.091) | 1.896 | 0.179 |
| | | | | | | Day | F(5,29.719) | 0 | 1 |
| | | | | | | Treatment*Day | F(5,29.091) | 2.183 | 0.083 |
| | | | | Pairwise (emmeans posthoc) | | | | | |
| | | | | **mean inoculated group** | **mean depleted group** | | | | |
| | | | 1 | 0.35 | 0.65 | Treatment | t-ratio | 1.814 | 0.08 |
| | | | 2 | 0.278 | 0.722 | Treatment | t-ratio | 2.688 | **0.012** |
| | | | 3 | 0.436 | 0.564 | Treatment | t-ratio | 0.889 | 0.381 |
| | | | 4 | 0.488 | 0.512 | Treatment | t-ratio | 0.162 | 0.872 |
| | | | 5 | 0.561 | 0.439 | Treatment | t-ratio | -0.848 | 0.403 |
| | | | 6 | 0.562 | 0.438 | Treatment | t-ratio | -0.873 | 0.39 |
| 6A | *L. melliventris* | Cumulative proportion bees foraging per treatment group | All | Cox Proportional Hazards | | Treatment | z | -0.435 | 0.664 |
| | | | | **mean inoculated group** | **mean depleted group** | | | | |
| | | | 1 | 0.034 | 0.05 | | | | |
| | | | 2 | 0.074 | 0.096 | | | | |
| | | | 3 | 0.208 | 0.217 | | | | |
| | | | 4 | 0.28 | 0.255 | | | | |
| | | | 5 | 0.326 | 0.32 | | | | |
| | | | 6 | 0.347 | 0.346 | | | | |
| 6B | *L. melliventris* | Proportion foraging events per treatment group | All | Linear Mixed Effects Model | | Treatment | F(1,31.021) | 0.351 | 0.558 |
| | | | | | | Day | F(5,31.443) | 0 | 1 |
| | | | | | | Treatment*Day | F(5,31.021) | 4.169 | **0.005** |
| | | | | Pairwise (emmeans posthoc) | | | | | |
| | | | | **mean inoculated group** | **mean depleted group** | | | | |

| | | | | | | | | | |
|---|---|---|---|---|---|---|---|---|---|
| | | | 1 | 0.324 | 0.676 | Treatment | t-ratio | 2.96 | **0.006** |
| | | | 2 | 0.305 | 0.695 | Treatment | t-ratio | 2.833 | **0.008** |
| | | | 3 | 0.539 | 0.461 | Treatment | t-ratio | -0.651 | 0.52 |
| | | | 4 | 0.585 | 0.415 | Treatment | t-ratio | -1.43 | 0.163 |
| | | | 5 | 0.543 | 0.457 | Treatment | t-ratio | -0.719 | 0.478 |
| | | | 6 | 0.571 | 0.429 | Treatment | t-ratio | -1.193 | 0.242 |
| 6C | *L. melliventris* | Number of foragers per treatment group | All | Linear Mixed Effects Model | | Treatment | F(1,33) | 0.079 | 0.781 |
| | | | | | | Day | F(5,33) | 18.926 | **<0.001** |
| | | | | | | Treatment*Day | F(5,33) | 0.132 | 0.984 |
| | | | | Pairwise (emmeans posthoc) | | | | | |
| | | | | **mean inoculated group** | **mean depleted group** | | | | |
| | | | 1 | 2 | 3 | Treatment | t-ratio | 0.344 | 0.733 |
| | | | 2 | 3.75 | 4.75 | Treatment | t-ratio | 0.344 | 0.733 |
| | | | 3 | 11.8 | 12 | Treatment | t-ratio | 0.086 | 0.932 |
| | | | 4 | 15 | 13.2 | Treatment | t-ratio | -0.601 | 0.552 |
| | | | 5 | 16.8 | 17.5 | Treatment | t-ratio | 0.258 | 0.798 |
| | | | 6 | 16.5 | 17.2 | Treatment | t-ratio | 0.258 | 0.798 |
| 6D | *L. melliventris* | Proportion foraging events per individual | All | Generalized Linear Mixed Effects Model | | Treatment | $\chi^2$ | 0.141 | 0.707 |
| | | | | | | Day | $\chi^2$ | 334.831 | **<0.001** |
| | | | | | | Treatment*Day | $\chi^2$ | 18.336 | **0.003** |
| | | | | Pairwise (emmeans posthoc) | | | | | |
| | | | | **mean inoculated group** | **mean depleted group** | | | | |
| | | | 1 | 0.162 | 0.225 | Treatment | z-ratio | 1.459 | 0.145 |
| | | | 2 | 0.061 | 0.11 | Treatment | z-ratio | 3.812 | **<0.001** |
| | | | 3 | 0.046 | 0.038 | Treatment | z-ratio | -0.096 | 0.923 |
| | | | 4 | 0.039 | 0.031 | Treatment | z-ratio | -0.076 | 0.94 |
| | | | 5 | 0.032 | 0.026 | Treatment | z-ratio | -0.096 | 0.924 |
| | | | 6 | 0.035 | 0.025 | Treatment | z-ratio | -0.495 | 0.62 |
| 6E | *L. melliventris* | Proportion of elite foragers per treatment group | All | Linear Mixed Effects Model | | Treatment | F(1,31.021) | 0.0006 | 0.981 |
| | | | | | | Day | F(5,31.443) | 0 | 1 |
| | | | | | | Treatment*Day | F(5,31.021) | 3.961 | **0.007** |
| | | | | Pairwise (emmeans posthoc) | | | | | |
| | | | | **mean inoculated group** | **mean depleted group** | | | | |
| | | | 1 | 0.292 | 0.708 | Treatment | t-ratio | 2.306 | **0.028** |
| | | | 2 | 0.194 | 0.806 | Treatment | t-ratio | 2.929 | **0.006** |
| | | | 3 | 0.637 | 0.363 | Treatment | t-ratio | -1.515 | 0.14 |
| | | | 4 | 0.61 | 0.39 | Treatment | t-ratio | -1.222 | 0.231 |
| | | | 5 | 0.582 | 0.418 | Treatment | t-ratio | -0.907 | 0.371 |
| | | | 6 | 0.603 | 0.397 | Treatment | t-ratio | -1.139 | 0.263 |

**Supplementary Table 5: Statistics for each single microbe inoculation behavioral analyses requiring pairwise comparisons depicted in Figures 4-6.** Linear mixed effects model results for most behavioral measures per single microbe inoculation experiment, with inoculation treatment and day as main factors and replicate as a random factor, followed by pairwise comparison results between inoculation treatments on each day. Generalized linear mixed effects model with log-normal distribution results for proportion foraging events per individual for each single microbe inoculation experiment, with inoculation treatment and day as main factors, and replicate and individual as random factors, followed by pairwise comparison results between inoculation treatments on each day. n = 4 colonies.

| Feature | Description |
|---|---|
| First Y | Barcode y-coordinate when the bee appears. |
| Last Y | Barcode y-coordinate when the bee disappears. |
| Horizontal displacement | Sum of horizontal barcode displacements. |
| Vertical displacement | Sum of vertical barcode displacements. |
| Duration | Duration of the pass. |
| Distance traveled | Sum of Euclidean distances moved. |
| Rotation | Sum of turning angles. |

**Supplementary Table 6:** Features calculated by the flight activity detector. Note that the openings through which bees can enter and exit the entrance monitor enclosure are located at the top and bottom edge of the recorded video. Further note that displacements and rotations are signed values.

| Pass class | Sensitivity | Positive predictive value | $F_1$ score |
|---|---|---|---|
| Incoming | 0.96 | 0.88 | 0.92 |
| Outgoing | 0.91 | 0.79 | 0.85 |

**Supplementary Table 7:** Flight activity detector performance.

### Supplementary Methods

#### Flight activity

##### Entrance monitor

Flight activity was recorded with an improved version of the entrance monitor described in [70]. Improvements aimed to make the entrance monitor easier to traverse by the bees, increase bCode detection rate and recording frame rate, and make the entrance monitor enclosure cheaper and easier to manufacture. The changes we made to achieve these goals are described below.

###### Enclosure

When passing through the entrance monitor enclosure, bees need to traverse a maze. This maze slows them down so they can be recorded multiple times while exiting or returning to the hive. The longer dimension of this maze was shortened to approximately half its original size. In addition, we removed two of the three inner walls. These changes helped the bees navigate the maze and made it less likely that they considered it part of their hive and congregated in it. The roof of the maze was changed to glass with an antireflective coating. This coating eliminated reflections of the camera and other enclosure components on the glass and thus helped to increase the barcode detection rate.

To manufacture the enclosure, we created a three-dimensional model of it (Supplementary Fig. 3) in Onshape (PTC Inc., Boston, Massachusetts, USA), an online computer-aided design software system. This model was printed on a Form 2 printer (Formlabs, Somerville, Massachusetts, USA), using Clear Resin (Formlabs, Somerville, Massachusetts, USA). Clear Resin cures to optical translucency, which reduced motion blur by permitting more light to pass through the enclosure walls than the opaque material we used before. After printing, the enclosure was cleaned in a Form Wash (Formlabs, Somerville, Massachusetts, USA) and cured in a Form Cure (Formlabs, Somerville, Massachusetts, USA), using manufacturer-recommended settings.

**Camera**

The entrance monitor camera was upgraded to a Raspberry Pi camera module v2.1 (Raspberry Pi Ltd, Cambridge, UK). The higher resolution of this camera made it possible to use pixel binning to record the maze at 1640 x 1232 pixels. This improved the camera's performance under low-light conditions and led to bigger barcodes in the recorded footage, which contributed to the higher barcode detection rate. For data storage, we switched from capturing images in burst mode to recording video. This enabled the camera to automatically adjust to changes in illumination. It also reduced the amount of data and made data storage more time-efficient. These changes allowed us to step up the recording frame rate to 10 Hz, a five-fold increase that enabled us to capture each bee multiple times despite them passing faster through the shorter, simplified maze.

**Performance**

To obtain an estimate of the entrance monitor identification rate, we manually annotated all bees that were not automatically identified in 17,125 images that were randomly sampled from entrance monitor videos recorded during an unrelated experiment. Of the bees with a barcode that was at least partially visible, 88.6 ± 26.1% (mean standard ± deviation) were automatically identified. Out of all bees in an image, including bees with a barcode that was not visible, 63.5 ± 36.9% were identified. The most common reasons for a bee not getting identified were that she walked on the roof of the maze (58.4%) or had lost her barcode (17.1%).

#### Hive exits and returns

To detect flight activity, we developed a new detector for hive exits and returns. Briefly, this detector groups temporally adjacent detections of each bee into passes. For each pass, it then computes features that describe the bee's movement through the maze. These features are fed to a random forest that classifies each pass as incoming, outgoing, or "other", whereby the "other" class accommodates bees entering and exiting the entrance monitor through the same opening (i.e., without traversing the maze). Details of the flight activity detector are described below.

##### Ground truth

To produce a data set for flight activity detector training and performance evaluation, we manually annotated all detected bees in 117 five min long video clips that were extracted from entrance monitor videos recorded during an unrelated experiment. Annotating a bee consisted of visually tracking her from the moment she appeared at one of the two openings of the entrance monitor until she disappeared, using custom video annotation software. The resulting trajectories were classified as incoming if the bee appeared at the outside-facing opening and disappeared through the hive-facing opening. Trajectories beginning at the hive-facing opening and ending at the outside-facing opening were classified as outgoing. All other trajectories were classified as "other".

Annotations were performed by a group of self-trained raters that had achieved a $F_1$ score of at least 0.9 in a one-time test that consisted of annotating a five min long gold-standard clip that had been annotated by an expert. The ground truth produced by these raters comprised 645,447 barcode detections in 9,481 trajectories, and was split into disjunct training and test sets consisting of 70% and 30% of the trajectories, respectively. The training set was subsampled to ensure that all three classes were represented equally, which reduced its size to 62% of the ground truth trajectories.

**Pass detection and classification**

Our flight activity detector first groups a bee's barcode detections that are at most $c$=30 s apart into a pass. The cutoff $c$ corresponds to the 99.9th percentile of the time between barcode detections in the training set, but was otherwise chosen arbitrarily. For each pass, the detector then computes the features listed in Supplementary Table 6 on the subset of successive barcode detections. Feature calculations were limited to this subset to ensure that pass features are not calculated across barcode detection gaps. Next, the detector predicts the pass class (i.e., whether it is an incoming, outgoing, or "other" pass), using a random forest consisting of 50 decision trees. This random forest was trained with default parameters on the training set trajectories, using the R package randomForest [71].

**Performance**

If the flight activity detector split an annotated pass into multiple detected passes, we counted only one of the detections as a positive. The remaining detections were considered to be false positives and thus decreased the detector's positive predictive value. Similarly, if a detected pass spanned multiple annotated passes, only one of these passes was considered to be detected. The other passes were treated as false negatives and thus decreased the detector's sensitivity. Finally, since the purpose of the flight activity detector is to identify hive exits and returns, its performance (Supplementary Table 7) on the "other" class was not evaluated and detections classified as "other" were ignored when calculating the performance on the incoming and outgoing class.
